# Biocatalytic Production of Galantamine in Yeast through Cytochrome P450 Optimization

**DOI:** 10.64898/2026.09.09.750345

**Authors:** Maxence Holtz, Samantha de Haan, Cecília Castellví Domingo, Niklas G. Madsen, María De Fátima Salazar, Frederik G. Hansson, Aafke C. A. van Aalst, Jonathan Asmund Arnesen, Christoph Crocoll, Simon d’Oelsnitz, Michael K. Jensen, Carlos G. Acevedo-Rocha

## Abstract

Galantamine is a pharmaceutically relevant *Amaryllidaceae* alkaloid used for the treatment of Alzheimer’s disease. Its structural complexity and low abundance in plants motivate the development of alternative manufacturing routes. Here, we established the first engineered yeast platform for the biocatalytic production of galantamine from 4OMe-norbelladine. Heterologous expression of the downstream pathway enzymes NtCYP96T6, NtNMT1, and NtAKR1 from *Narcissus cv. Tête-à-Tête* in *Saccharomyces cerevisiae* was complemented by optimization of cultivation temperature, carbon source, and medium pH enabling the first demonstration of galantamine production in yeast. Systematic screening of cytochrome P450 reductase and cytochrome b₅ partners to boost NtCYP96T6 activity led to the identification of a novel reductase mined from the *Narcissus pseudonarcissus* transcriptome, NpCPR, which supported the highest pathway flux and yielded 7.0 ± 0.6 mg/L galantamine, corresponding to a 7.6 % molar yield from 250 μM 4OMe-norbelladine and an approximately 173-fold improvement over the parental strain. We further exploited this yeast cell factory for the precursor-directed biosynthesis of 7F-galantamine, highlighting the potential of pathway enzyme promiscuity to access new-to-nature GAL analogues that may be challenging to produce through conventional chemical synthesis. Together, this work establishes a foundation for microbial galantamine production and biosynthetic diversification of its pharmaceutically relevant scaffold.

## Introduction

Galantamine (GAL) is an FDA-approved drug for the treatment of dementia and Alzheimer’s disease through a potent inhibitory activity against acetylcholine esterase. It belongs to the *Amaryllidaceae* alkaloids (AAs) family and it is produced by several plant species including *Galanthus nivalis* (common snowdrop) or *Narcissus pseudonarcissus* (wild daffodil)^1,2^. Although GAL is orally bioavailable and crosses the blood-brain barrier, its clinical benefit is moderate, and dose escalation is limited by peripheral cholinergic adverse effects^3,4^. Its relatively short elimination half-life of approximately 7-8 h and metabolism by CYP2D6 and CYP3A4 also leaves room for improvement of its pharmacokinetic profile and brain-to-blood ratio^5^. Continued interest in optimizing GAL-based therapeutics is illustrated by the recent FDA approval of the GAL prodrug benzgalantamine marketed as Zunveyl^6^.

The structure of GAL includes a strained tetracyclic framework and three stereo-centers making it highly challenging to produce at scale through total chemical synthesis^7^. As a result, these multi-step chemical synthesis processes are not economically competitive compared to extraction from native plants due to low overall yield. Plant extraction is therefore, as of now, the main supply source of GAL for pharmaceutical applications^8^. However, large-scale cultivation of GAL-producing plants is constrained by slow crop establishment, seasonal harvesting, and low GAL concentrations in fresh biomass (∼0.02-0.05 % w/w). Consequently, large quantities of plant biomass must be cultivated and processed, resulting in low productivity and contributing to historically high GAL prices of approximately US$ 40,000/kg^2,9^. While it is estimated that around 55 million people live with dementia as of 2024, the demand for GAL has been steadily increasing with a market valued at valued at US$ 1,000 million in 2023 and predicted to reach US$ 2,110 million by the end of 2030^10^. Therefore, there is a high need for efficient GAL supply chains industrially.

The GAL biosynthesis pathway was elucidated recently (**Fig. 1**)^11^. All AAs are made from 3,4-dihydroxybenzaldehyde (3,4-DHBA) (**1**) and tyramine (**2**). These two precursors are condensed through a Mannich reaction to form norcraugsodine (**3**) and then norbelladine (**4**) through the respective actions of norbelladine synthase (NBS) and norcraugsodine reductase (NR)^12^. Norbelladine is then 4’O-methylated by N4OMT to form 4OMe-norbelladine (**5**) which is the common precursor to all >650 AAs discovered so far^1^. This diversity is achieved by a phenol coupling reaction catalyzed by a cytochrome P450 monooxygenase (CYP) followed by action of tailoring downstream enzymes. In the case of GAL biosynthesis in *Narcissus cv. Tête-à-Tête*, it is NtCYP96T6 which performs a para’-ortho coupling leading to nornarwedine (**6**), which is then N-methylated by NtNMT1 to form narwedine (**7**). A final reduction step catalyzed by NtAKR1 leads to GAL (**9**)^13^.

**Figure 1:**
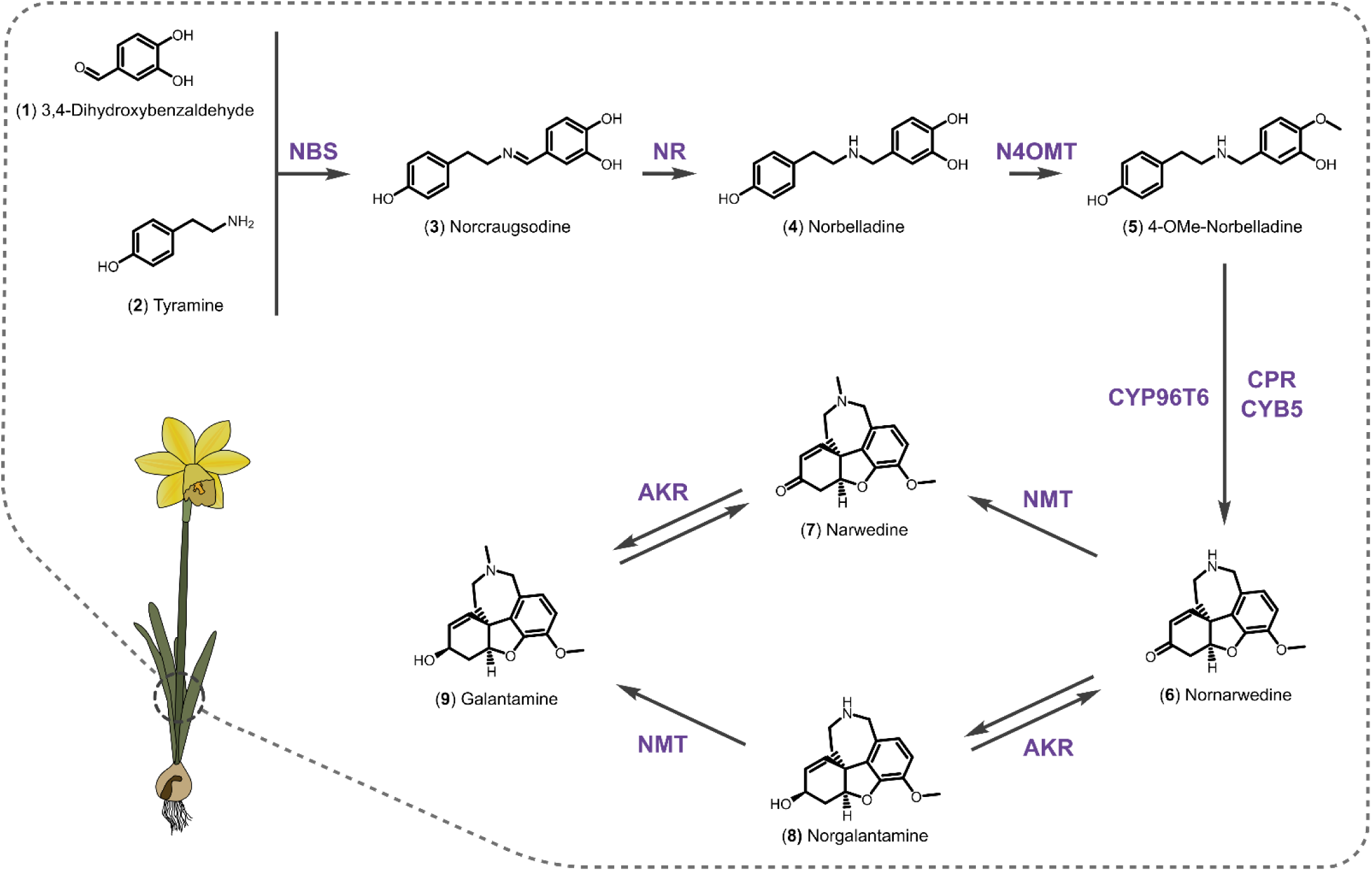
Biosynthetic pathway for galantamine production. Pathway precursors are tyramine and 3,4-dihydroxybenzaldehyde (3,4-DHBA) which are condensed and reduced to norbelladine by NBS and NR respectively. Norbelladine then gets O-methylated at C4 cyclized by a cytochrome P450 and N-methylated / reduced to yield galantamine (GAL). Abbreviations are: NBS, norbelladine synthase; NR, norcraugsodine reductase; N4OMT, norbelladine 4-O-methyltransferase; CYP96T6, cytochrome P450 96T6; CPR, cytochrome P450 reductase; CYB, cytochrome b5; NMT, N-methyltransferase; AKR, aldo-keto reductase.

The outlined limitations of plant extraction and total chemical synthesis is the motivation for the development of a heterologous microbial platform for GAL production. Such a platform could provide a potentially scalable and season-independent alternative supply route while exploiting enzyme selectivity to perform transformations that are difficult to achieve chemically. In particular, the P450-catalyzed oxidative phenol coupling reaction that introduces the correct stereochemistry at the tricyclic benzofuran core of GAL is an important bottleneck for synthetic chemistry^14^. In this study, we demonstrate for the first time the bioconversion of 4OMe-norbelladine to GAL in engineered *Saccharomyces cerevisiae* expressing NtCYP96T6, NtNMT1 and NtAKR1. We identified optimal redox partners for NtCYP96T6 and optimized cultivation conditions yielding a 173-fold improvement in GAL titer reaching 7.0 ± 0.6 mg/L (7.6 % molar yield) from 250 µM fed 4OMe-morbelladine. As an additional demonstration of the platform’s versatility, precursor-directed biosynthesis enabled the production of 7F-GAL that, to the best of our knowledge, have not previously been accessed through chemical synthesis. Together, these results establish a yeast cell factory for GAL biomanufacturing and derivatization.

## Results and Discussion

### Bioconversion of 4OMe-norbelladine to galantamine in yeast

To mitigate challenges related to chemical formation of the correct stereochemistry at the tricyclic benzofuran core of GAL, we aimed to construct an alternative route for a P450-catalyzed oxidative phenol coupling reaction to enable GAL production from 4OMe-norbelladine in *S. cerevisiae*. For this purpose, we initially cloned the downstream pathway module genes NtCYP96T6, NtNMT1 and NtAKR1 and integrated them into the genome of our base strain MIA-B0 generating strain yCCD45. This strain was grown in Synthetic Complete (SC) media supplemented with 2 % glucose with and without 250 µM 4OMe-norbelladine (**5**) for 144h at 30°C. The supernatants were subsequently analyzed by Liquid Chromatography-High Resolution Tandem Mass Spectrometry (LC-MS/MS) to assess the presence of AAs. While we could detect putative peaks matching the mass of nornawedine (**6**) ([M+H]^+^ m/z=272.1281), narwedine (**7**) ([M+H]^+^ m/z=286.1437) and norgalantamine (**8**) ([M+H]^+^ m/z=274.1437) when the pathway substrate 4OMe-norbelladine was fed, we could not detect any GAL being produced (**Supp. Fig. 1**). yCCD45 also produced another peak with the same mass as nornawedine (**6**) which we hypothesized to be noroxomaritidine, the product of para-para’ coupling of 4OMe-norbelladine (**Supp. Fig. 1**). This indicates some level of promiscuity of NtCYP96T6 in the yeast cell context which was also recently reported in tobacco leaf transient expression experiments^11^.

To improve pathway flux, we subjected yCCD45 to a small-scale optimization of bioconversion conditions by varying cultivation temperature, carbon source and pH. Production of narwedine (**7**) increased 5.4-fold by switching from 30°C to 25°C (**Fig. 2A**), potentially reflecting improved folding or stability of the heterologous pathway enzymes. Replacing 2% glucose with 2% trehalose resulted in a further 1.8-fold increase (**Fig. 2B**). Trehalose may support heterologous enzyme stability^15^ and, because it is assimilated more slowly than glucose, may also provide a carbon-feeding profile resembling fed-batch cultivation^16^. This has been previously observed during the optimization of complex plant alkaloid pathways in yeast^17,18^. The largest effect was obtained by increasing the medium pH from 5.5 to 7.0, which increased narwedine production by 25.0-fold (**Fig. 2C**). We hypothesized that this improvement resulted from enhanced transport of 4OMe-norbelladine into the yeast cells. To test this hypothesis, we functionalized in yeast a previously published 4OMe-norbelladine biosensor^19^ using our previously established yeast RamR sensor platform^20^. The intracellular biosensor signal was >3.7-fold higher at pH 7.0 than at pH 5.5 (**Supp. Fig. 2**), supporting increased intracellular availability and improved uptake of 4OMe-norbelladine under near-neutral conditions.

**Figure 2:**
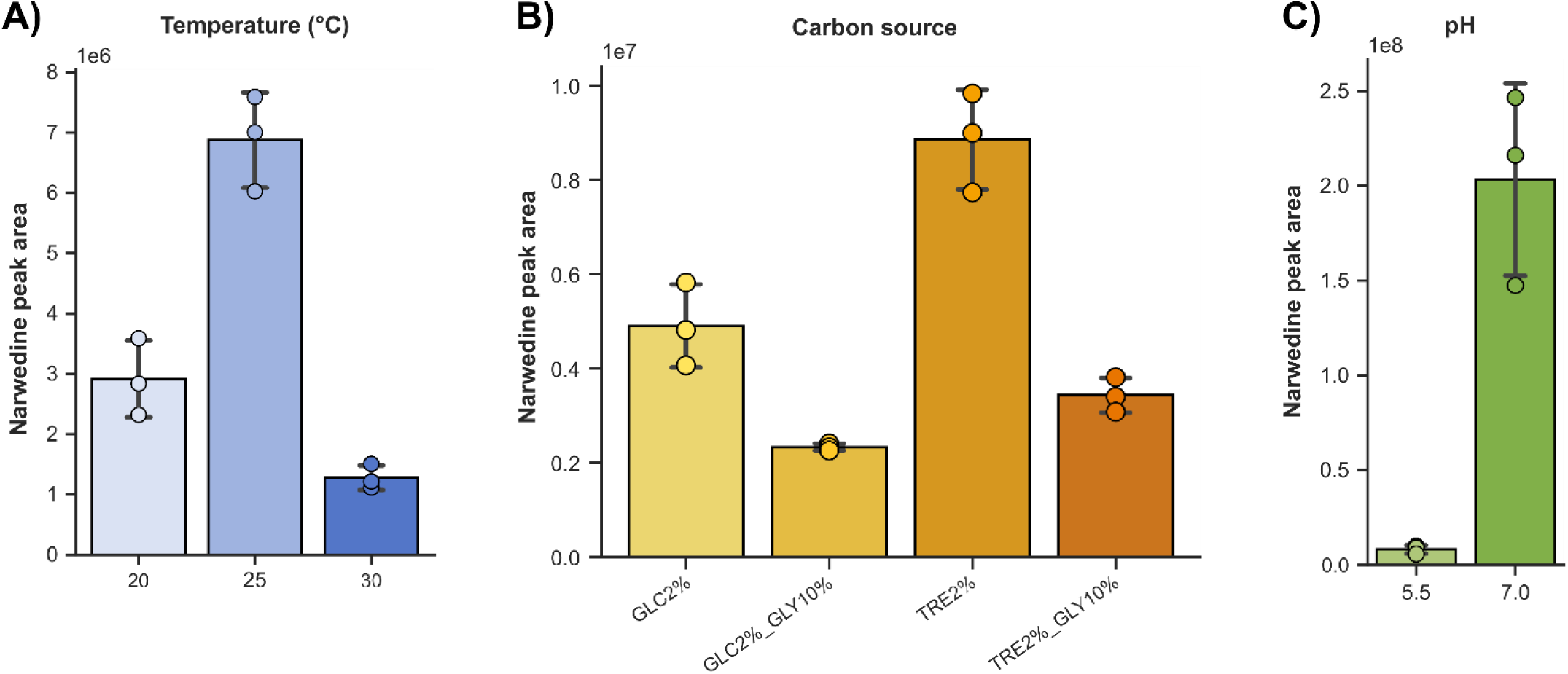
Optimization of cultivation conditions for narwedine production by yCCD45. Different temperatures **(A)**, carbon sources **(B)** and media pH **(C)** were tested for increased narwedine production by yCCD45 in synthetic complete media supplemented with 250 µM 4OMe-norbelladine. Production was carried out for 6 days and narwedine accumulation was measured with LC-MS/MS. Carbons source abbreviations are: GLC, glucose; TRE, trehalose; GLY, glycerol. Carbons source concentrations are indicated in % w/v.

These optimized conditions enabled the detection of a peak eluted at the same retention time and that had identical MS/MS fragmentation as authentic analytical standards of GAL (RT = 4.25 min) which was not present in the supernatant from either the parent MIA-B0 or yCCD45 without fed 4OMe-norbelladine. The GAL titer from yCCD45 in these optimized conditions was ∼ 41 µg/L. Our results demonstrate the functionality of NtCYP96T6, NtNMT1 and NtAKR1 in yeast representing the first reported production of GAL in engineered yeast (**Fig. 3, Supp. Fig. 3**). Interestingly, NtCYP96T6 appears capable, albeit inefficiently, of coupling with the native yeast cytochrome P450 reductase (CPR). This observation contrasts with the findings of Wu *et al.*, (2024) who reported that heterologous expression of this enzyme in yeast required co-expression of a plant-derived CPR to achieve detectable activity. This discrepancy may be explained by substantial differences in strain design and bioconversion conditions between the studies.

**Figure 3:**
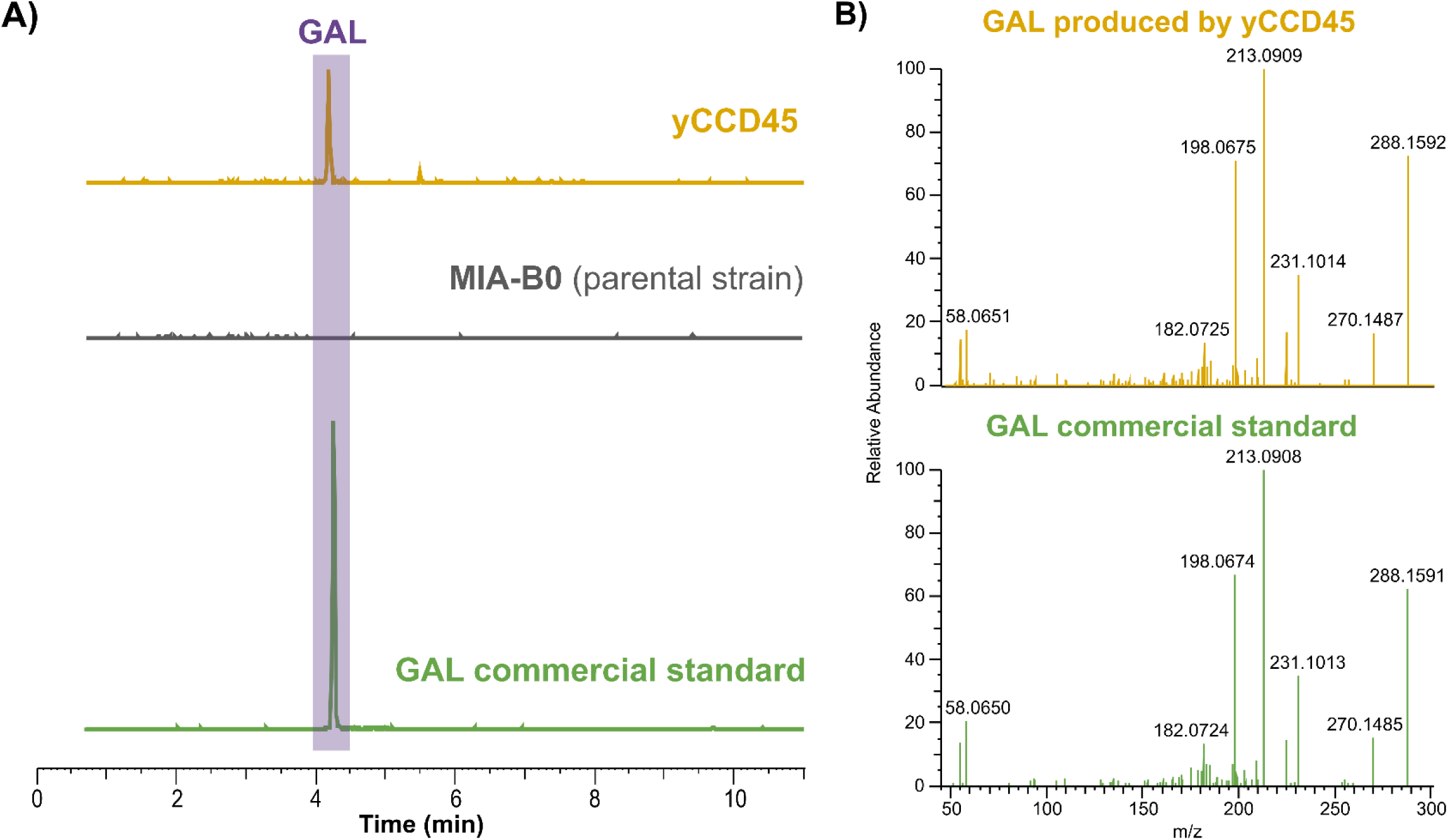
Proof of concept of GAL production by yCCD45 in optimized cultivation conditions. **A)** Detection of GAL produced by yCCD45 grown in SC 2% trehalose pH 7.0 supplemented with 250 µM 4OMe-norbelladine at 25°C (EIC m/z = 288.1594, 5 ppm mass resolution). **B)** Representative MS/MS spectra of GAL produced by yCCD45 compared to an authentic commercial standard.

### Redox partner screening enhances NtCYP96T6 activity and galantamine production in yeast

As most of the downstream pathway substrate 4OMe-norbelladine remained unconverted, we next sought to identify heterologous CPR and cytochrome b_5_ (CYB5) partners that could enhance NtCYP96T6 catalysis in yeast cells, given the well-established dependence of plant P450 enzymes on compatible electron transfer partners^21^. Plant CYPs are bound to the endoplasmic reticulum by their N-terminal domain and need to receive electrons originating from NADPH oxidation through protein-protein interaction with the CPR (**Fig. 4A**). CYB5 can act as an extra redox partner, providing electrons through NADH oxidation while stabilizing and allosterically modulating the CYP-CPR complex^21^.

**Figure 4:**
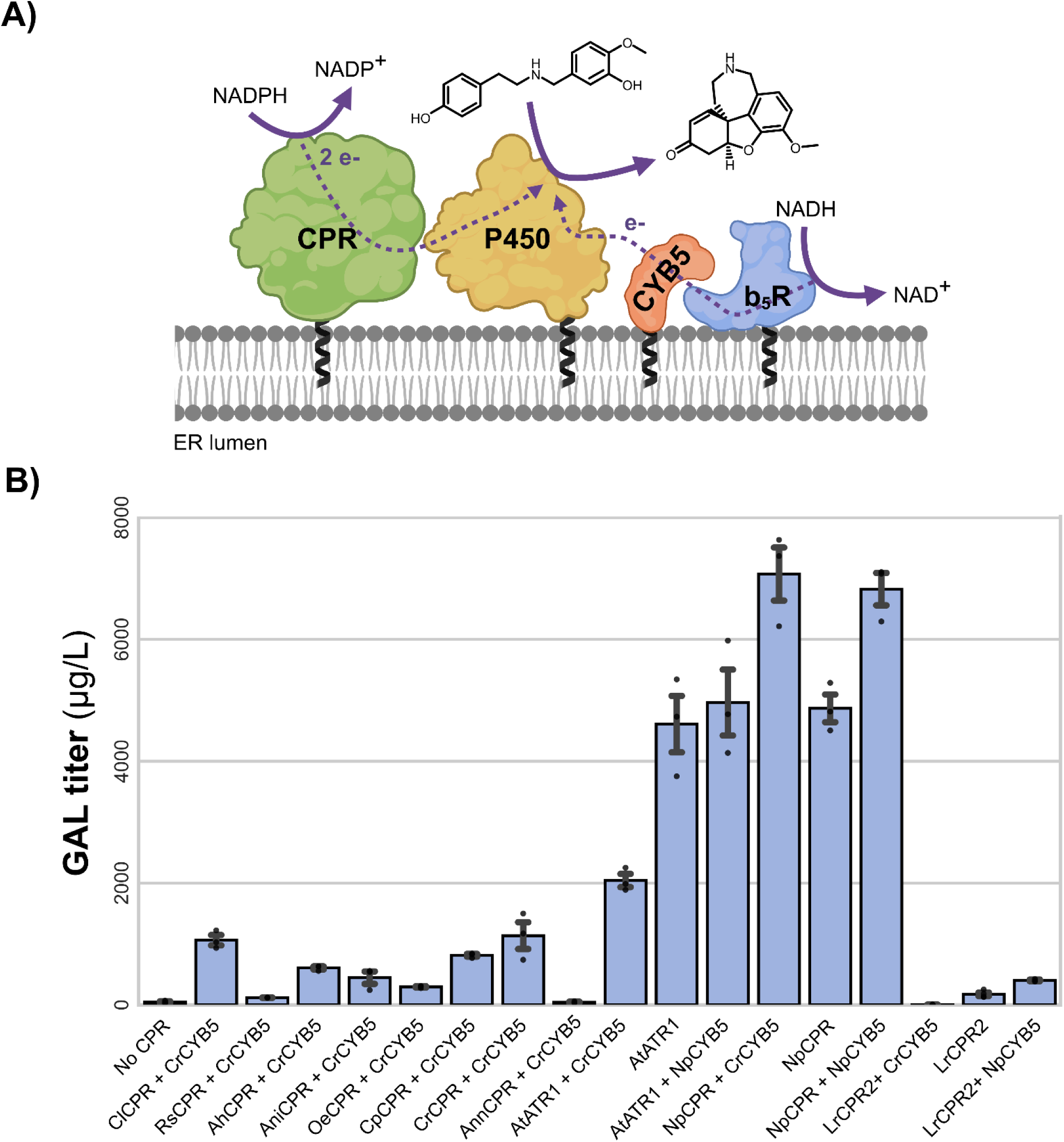
Optimization of NtCYP96T6 redox partners enhances galantamine production in yeast. **A)** Schematic of electron transfer from CPR to the ER-bound plant P450 and the auxiliary electron-transfer pathway involving CYB5 and NADH-cytochrome *b*₅ reductase (b₅R). Solid arrows indicate cofactor or substrate oxidation, and dashed arrows indicate electron transfer between proteins. **(B)** Galantamine production by *S. cerevisiae* strains expressing NtCYP96T6, NtNMT1 and NtAKR1 with the indicated CPR and CYB5. combinations. Strains were cultivated with 250 µM 4OMe-norbelladine in synthetic complete medium containing 2% trehalose at pH 7.0 and 25 °C for 144 h. Galantamine titers were quantified by LC–MS/MS. Abbreviations are: Cl, *Catharanthus longifolius*; Rs, *Rauvolfia serpentina*; Ah, *Amsonia hubrichtii*; Ani, *Aspergillus niger*; Oe, *Olea europaea*; Cp, *Chrysomela populi*; Ann, *Artemisia annua*; At, *Arabidopsis thaliana*; Cr, *Catharanthus roseus*; Lr, *Lycoris radiata*; Np, *Narcissus pseudonarcissus*. Measurements for each condition represent the average of three biological replicates. Error bars represent S.D.+/−the mean.

We started from a panel of nine plant and fungal CPRs coming from previous work in our laboratory^22^ which were all co-expressed with CrCYB5 (from *Catharantus roseus*) (**Supp. Table 1**). We also included the top performing candidate from *Lycoris radiata* LrCPR2 identified by Wu *et al.*, (2024). Finally, because a transcriptome for *Narcissus cv. Tête-à-Tête*, the source of NtCYP96T6, was not available at the time, we mined the transcriptome of the closely related daffodil *Narcissus pseudonarcissus*^23^ to identify candidate CPR and CYB5 redox partners. While these proteins cannot be described as strictly co-evolved with NtCYP96T6, their identification from a related *Narcissus* species supports their use as plausible redox partners in yeast. Using the *Arabidopsis thaliana* AtATR1 as query, we found a sequence with 67 % identity that we tentatively name NpCPR.

All these eleven CPR candidates were genome integrated together with CrCYB5 in yCCD45 and the resulting strains were tested for their GAL production capabilities when 250 µM 4OMe-norbelladine was fed in the optimal bioconversion conditions (SC 2% trehalose, pH 7, 25°C). Nearly all the tested heterologous CPRs improved GAL production indicating that coupling with the native yeast CPR was not enough to support efficient NtCYP96T6 activity (**Fig. 4B**). The highest production was achieved with our newly discovered NpCPR reaching 7.0 ± 0.6 mg/L GAL (7.6 % molar yield), representing a ∼173-fold increase in titer compared to the parental strain yCCD45 and ∼29-fold increase over the published LrCPR2 without heterologous CYB5 under identical assay conditions ^24^. The magnitude of this improvement points to a particularly productive interaction between NpCPR and NtCYP96T6, potentially involving more efficient electron transfer through improved protein compatibility, or greater functional expression in yeast.

To further investigate the effect of CYB5 on NtCYP96T6 activity, we tested the top two CPRs (NpCPR and AtATR1) and the published LrCPR2 candidate with and without CrCYB5. We also included a new CYB5 candidate from *N. pseudonarcissus* mined from the transcriptome named NpCYB5 that had 79% identity with CrCYB5. The highest production was again obtained with NpCPR and the titer was significantly higher when either CrCYB5 or NpCYB5 were expressed indicating that CYB5 has an active role promoting catalysis in the yeast system. However, the difference in product formation between CrCYB5 and NpCYB5 was not statistically significant.

### Mitigation attempts for improved narwedine reduction

Although redox partner optimization improved GAL production, several high-performing strains still accumulated large amounts of narwedine (**Supp. Fig. 4**). While the absence of an authentic narwedine standard prevented absolute quantification, the strong LC-MS signal suggests that the terminal reduction of narwedine to GAL by NtAKR1 became limiting once flux through the P450-catalyzed step was increased. This interpretation is consistent with the recent characterization of GAL biosynthetic enzymes in *Leucojum aestivum*, where AKRs were shown to reversibly interconvert both nornarwedine/norgalanthamine and narwedine/galanthamine, with individual homologs displaying different directional preferences^25^. LaAKR3 favored the reductive direction, whereas LaAKR1 favored oxidation, highlighting that AKR choice can strongly influence whether flux is pulled toward or away from GAL formation. To relieve this bottleneck, we overexpressed NtNMT1 and NtAKR1 from high-copy 2µ plasmids in the high producer yMHO118 generating strain yMHO247. However, this did not further improve conversion of narwedine to GAL (**Supp. Fig. 4**). These results suggest that enzyme abundance alone is not sufficient to overcome the terminal reductase bottleneck. Further improvements in GAL titer may therefore require screening additional AKR homologs from GAL-producing *Amaryllidaceae* species, particularly reductive-biased enzymes, or engineering NtAKR1 to enhance productive narwedine recognition and reduction while minimizing competing substrate utilization and alternative reaction outcomes.

Further analysis of the untargeted metabolomics data also led to the identification of features matching the theoretical masses of norlycoramine ([M+H]+ m/z=276.1594) and lycoramine ([M+H]+ m/z=290.1751), the reduced derivatives of norgalantamine and galantamine respectively (**Supp. Fig 5**). Both compounds have previously been reported as naturally occurring *Amaryllidaceae* plants^25^. These putative products suggest further reduction of the GAL scaffold by NtAKR1 or endogenous yeast reductases, although further confirmation with authentic standards would be required.

**Figure 5:**
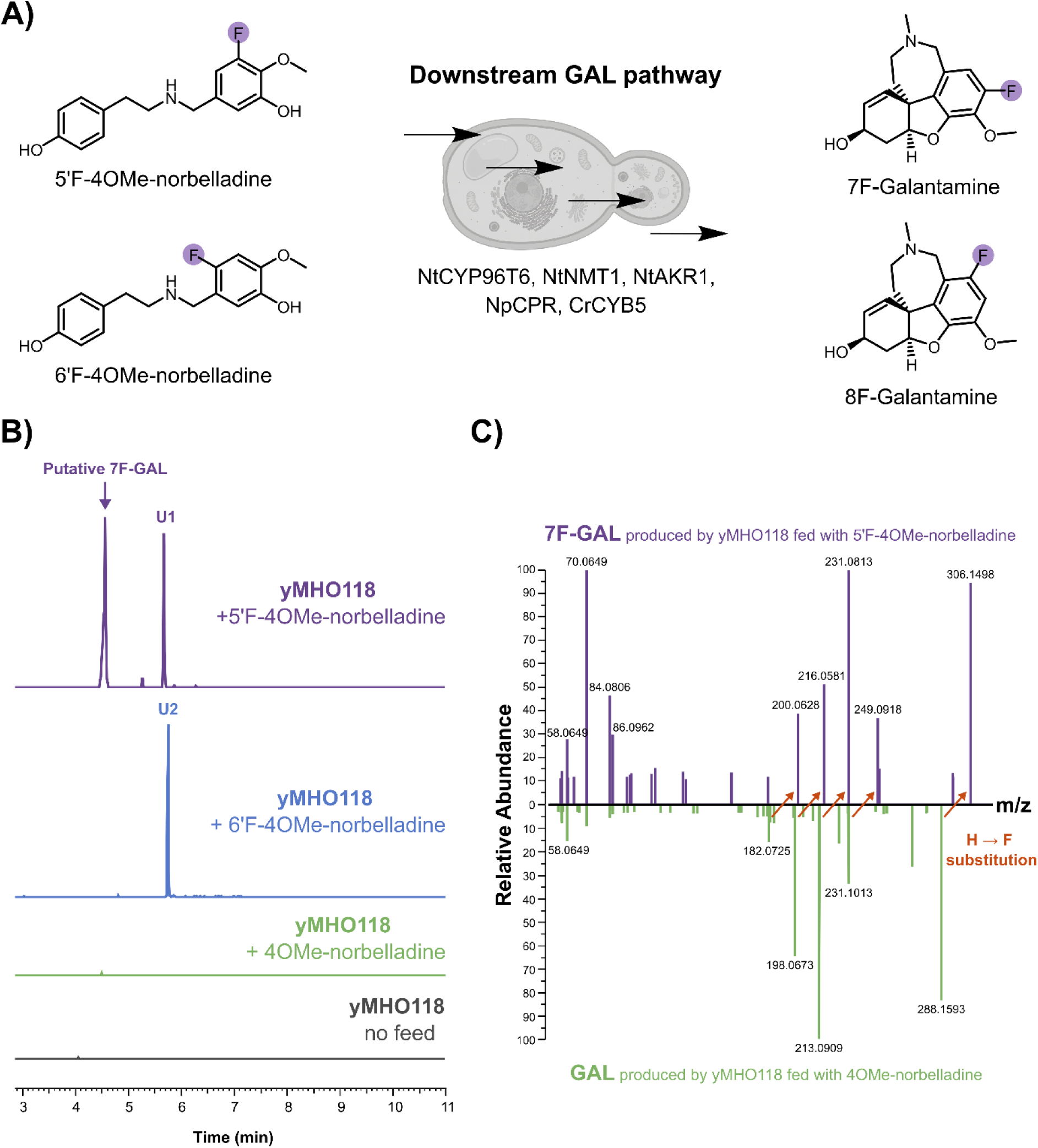
Precursor-directed biosynthesis of fluorinated galantamine derivatives in yeast. **A)** Experimental setup for bioconversion of 5′-fluoro- and 6′-fluoro-4OMe-norbelladine by engineered strain yMHO118 into the corresponding fluorinated galantamine derivatives. **B)** Monofluorinated GAL production from yMHO118 cultures supplemented with 5′-fluoro-4OMe-norbelladine (purple), 6′-fluoro-4OMe-norbelladine (blue), nonfluorinated 4OMe-norbelladine (green), or no precursor (gray) (EIC m/z = 306.1500, 5 ppm mass resolution). Pathway precursors were supplemented at 250 µM and strains were cultivated in the identified optimized conditions (SC 2 % trehalose, pH 7.0, 25°C). U1 and U2 indicate unidentified metabolites **(C)** Mirror MS/MS comparison of putative 7-F-GAL produced from 5′-fluoro-4OMe-norbelladine and GAL produced from nonfluorinated 4OMe-norbelladine. Orange arrows indicate the +17.9906 m/z shift expected upon H-to-F substitution in corresponding fluorine-retaining product ions described in **Supp.** Fig. 3.

### Precursor-directed biosynthesis of fluorinated galantamine derivatives

Having established a functional GAL pathway in yeast, we next investigated whether the platform could accept 4OMe-norbelladine analogs and thereby generate new-to-nature GAL derivatives. Fluorination was selected as a proof of concept because strategic incorporation of fluorine can modify the metabolic stability, membrane permeability, and pharmacokinetic properties of small molecules while introducing limited steric perturbation^26,27^. We specifically targeted the two aromatic positions adjacent to the methoxy substituent of the GAL scaffold by feeding the corresponding monofluorinated 4OMe-norbelladine analogs, herein designated 5’-fluoro-4OMe-norbelladine and 6’-fluoro-4OMe-norbelladine, to yMHO118 in the optimized GAL production conditions (**Fig. 5A**). These cultures were analyzed by LC-MS/MS, and we observed a peak that matched the exact mass of 7-fluorogalantamine (7F-GAL, m/z=306.1500) in the samples fed with 5’-fluoro-4OMe-norbelladine (**Fig. 5B**). The MS/MS fragmentation for that peak (RT = 4.50 min) showed a mass addition of 17.99 Da, coherent with the substitution of a hydrogen atom with fluorine on the aromatic ring of the molecule (**Fig. 5C**, **Supp. Fig. 3**). Together, the accurate mass and fragmentation pattern support the putative assignment of this product as 7F-GAL. In contrast, supplementation with 6′-fluoro-4OMe-norbelladine did not produce a detectable peak consistent with 8-fluorogalantamine (8F-GAL) under the conditions tested. However, we detected a feature matching the expected mass and fragmentation pattern of 8-fluoronarwedine (**Supp. Fig. 6**), indicating that the fluorinated precursor underwent NtCYP96T6-catalyzed oxidative phenol coupling but was not efficiently converted by NtAKR1 to 8F-GAL. Several additional unidentified metabolites were detected following supplementation with the custom-synthesized fluorinated precursors. These signals may represent alternative pathway products or metabolic shunts but could also originate from impurities in the chemically synthesized precursor preparations.

The detection of putative 7F-GAL and 8-fluoronarwedine demonstrates that NtCYP96T6 tolerates fluorine substitution corresponding to both the C7 and C8 positions of the GAL scaffold. To the best of our knowledge, 7F-GAL has not previously been produced, and GAL derivatives bearing substitutions at C7 have not been reported. The chemical synthesis of racemic 8-F-GAL and its separation into the corresponding enantiomers were previously reported, although quantitative pharmacological data were not provided^28^. Overall, these results demonstrate that our engineered yeast platform can exploit the substrate promiscuity of the pathway enzymes to diversify the GAL scaffold through precursor-directed biosynthesis. The apparent acceptance of substitution at C7 is particularly promising because feeding precursors bearing larger halogens, such as bromine, could provide functional handles for subsequent cross-coupling reactions^26^. Combining enzymatic scaffold construction with downstream chemical derivatization could therefore enable access to a broader and previously unexplored collection of GAL analogues, establishing this yeast platform as a versatile foundation for future GAL diversification and drug discovery.

## Conclusions

In this study, we established the first engineered yeast platform for the biocatalytic production of galantamine from 4OMe-norbelladine. Functional reconstruction of the downstream pathway required the coordinated expression of NtCYP96T6, NtNMT1, and NtAKR1 in *Saccharomyces cerevisiae*. Optimization of the cultivation conditions enabled initial galantamine production, while systematic screening of heterologous redox partners identified NpCPR as a particularly effective reductase for NtCYP96T6 in yeast which when expressed with a plant cytochrome b_5_ resulted in a GAL titer of 7.0 ± 0.6 mg/L and a molar yield of 7.6% from the pathway precursor. This represented an approximately 173-fold improvement over the strain relying on the endogenous yeast redox system. Metabolomic analysis revealed the accumulation of pathway intermediates and several putative side products. In particular, narwedine accumulation suggests that the terminal reduction step represents a downstream bottleneck and motivates future screening of NtAKR1 homologs or engineering variants with increased specificity toward the reduction reaction. Extending beyond GAL biomanufacturing, we demonstrate precursor-directed biosynthesis of fluorinated GAL using an engineered yeast cell factory, highlighting a microbial route for expanding the chemical diversity of this pharmaceutically relevant scaffold. Together, these advances establish yeast as a versatile platform for the biomanufacturing and diversification of GAL and related *Amaryllidaceae* alkaloids.

## Materials & Methods

### Chemical standards

All chemical standards had a purity of 95% or higher. GAL was purchased from Sigma Aldrich (product number 345670). 4OMe-norbelladine was purchased from Toronto Research Chemicals (product number H948930). Fluorinated 4OMe-norbelladine derivatives were obtained through custom chemical synthesis (WuXi AppTech, China).

### Plasmid and Yeast Strain Construction

All plasmids used in this study were assembled using NEBuilder HiFi assembly (New England Biolabs) or using the Yeast MoClo (modular cloning) toolkit^29^. Assembly reactions were transformed into *E. coli* DH5α competent cells and plated on Luria–Bertani agar containing 100 μg.ml^−1^ ampicillin, 50 μg.ml^−1^ kanamycin or 25 μg.ml^−1^ chloramphenicol and grown at 37 °C overnight. Colonies were picked, grown overnight in 5 mL LB with appropriate antibiotics; the next day plasmid DNA was extracted (Nucleospin Plasmid, Macherey-Nagel) and sent for Sanger sequencing (Eurofins Genomics). The full list of gene sequences and plasmids used in this study are available respectively in **Supp. Table 1** and **Supp. Table 2**.

MoClo plasmids followed the hierarchical DNA construction workflow as described in the original Yeast MoClo publication^29^. Coding sequences of pathway genes were codon-optimized for *S. cerevisiae* and synthesized by Integrated DNA Technologies with relevant flanking type IIS restriction enzyme sites and overhangs. These synthesized fragments were assembled into the Level 0 entry vector pYTK001 using the NEBridge Golden Gate Assembly Kit (BsmBI-v2) (New England Biolabs) following the manufacturer’s protocol. Level 1 plasmids were built by combining a promoter, CDS and terminator part into pre-assembled pMHGG1_Vec suite containing a yeast URA selection marker, an *E. coli* ampicillin resistance gene, a CEN/ARS yeast origin of replication, connectors determining position of transcription unit in pathway and a GFP dropout cassette for visual screening of correct assembly in *E. coli* (NEBridge Golden Gate Assembly Kit BsaI-v2). Finally, these Level 1 transcription unit plasmids were combined with BsmbI-v2 into Level 2 EasyClone-MarkerFree CRISPR-Cas9 integration compatible vector from our previous work^17,30,31^.

All the integrative plasmids generated in this study were linearized using NotI (New England Biolabs) and the DNA fragments were transformed in yeast together with relevant gRNA plasmid from EasyClone-MarkerFree^30,31^ using standard lithium acetate methods^32^. The integration of heterologous genes was verified as previously described^30,31^.

### Media and yeast cultivation

All yeast strains used and constructed in this study (**Supp. Table 3**) are based on CEN.PK2-1C. For routine propagation and transformation purposes, yeast strains were grown at 30°C in YPD (yeast extract 1 % w/v, peptone 2 % w/v and glucose 2 % w/v). Unless specified otherwise, strains were grown in SC medium supplemented with 250 µM 4OMe-norbelladine to test for GAL production. The same was applied for GAL fluorination experiments using fluorinated 4OMe-norbelladine derivatives. Strains were inoculated from plate in 2 mL of YPD in 15 mL preculture tubes and incubated overnight at 30°C and 300 r.p.m shaking. The following day, the OD_600_ of each preculture was measured, and the cultures were normalized to an OD_600_ of 10 in the corresponding production medium. Aliquots of 300 µL were transferred to a 96-deep-well plate for production. All cultivations were run as biological triplicates originating from different colonies. Plates were incubated at specified temperature for 144 h with shaking at 300 r.p.m. Following 144 h, 100 μL of each sample was filtered through a filter plate (PALL, AcroPrep Advance, 0.2-μm Supor membrane for medium/water) by centrifugation at 2,200 g for 1.5 min. Samples were stored at −20°C prior to injection.

### Metabolite analysis by liquid chromatography-high resolution tandem mass spectrometry

Samples generated in this study were analyzed using an untargeted metabolomics system constituted of a Vanquish Duo UHPLC binary system (Thermo Fisher Scientific, USA) connected to an Orbitrap ID-X Tribrid mass spectrometer (Thermo Fisher Scientific. USA). Chromatographic separation was achieved under reverse-phase conditions as previously described^22,33^. The MS measurements were performed in positive-heated electrospray ionization mode with a voltage of 3,500 V acquiring the full MS/MS spectra (data-dependent acquisition-driven MS/MS) in the mass range of 70–1,000 Da. The following data-dependent acquisition settings were used: automatic gain control target value of 4 × 10^5^ for full-scan MS and 5 × 10^4^ for the MS/MS spectral acquisition and a mass resolution of 120,000 for full-scan MS and 30,000 for MS/MS events. Precursor ions were fragmented by stepped high-energy collision dissociation using collision energies of 20, 40 and 50. Because authentic commercial standards were unavailable for some GAL pathway intermediates or fluorinated GAL, their identities were assigned putatively based on exact mass, retention time, and characteristic MS/MS spectral shifts relative to the corresponding GAL standard fragmentation.

### Untargeted metabolomics data processing

LC-MS data were qualitatively inspected using the FreeStyle Software (Thermo Fischer Scientific, USA). The peaks integration was done using Thermo Xcalibur Quant Browser Software (Thermo Fisher Scientific, USA) using 5-ppm mass tolerance. The integrated peaks were also manually inspected to ensure correct peak assignment and integration. For shunt product identification the raw files were processed using MZmine 2.53^34,35^ to generate a table of metabolic features. The generated feature files were used as input for GNPS FBMN (Feature-Based Molecular Networking) which is one of the most significant and often used tools for molecular networking, annotation, and visualization in the field of metabolomics^36,37^. Features of interest found in clusters containing known AAs were computed using the SIRIUS executable for formula and structural predictions^38^.

### RamR biosensor assay for 4OMe-norbelladine

Yeast strains carrying the RamR 4NB2 biosensing platform^19,20^ were grown overnight in SC-URA (plasmid based RamR-4NB2 expression) at 30 °C and 250 r.p.m. The following day, the precultures were diluted 1:25 in fresh SC-URA at pH 5.5 or 7.0. From these cell suspensions, 180 µL was transferred per well in a 96-deep-well plate. The plate was incubated for 2 h at 30 °C and 250 r.p.m before inducing the biosensors with 4OMe-norbelladine. 20 µL of ligand in SC media was used as inducer bringing the DMSO concentration to 1 % v/v for any assay. Following additional 6 h of incubation at 30 °C and 250 r.p.m, 50 µL of cells were mixed with 100 µL of PBS in a microtiter plate and fluorescence was analyzed using a NovoCyte Quanteon (Agilent) flow cytometer. For each condition, three biological replicates were analyzed with a threshold of 20,000 events per replicate. The cells were gated for singlets in the exponential phase and the median fluorescence intensity (MFI) for the GFP signal (FITC-H channel) was determined using FlowJo v10.8 Software (BD Life Sciences).

## Supporting information

Holtz et al, 2026_Supplementary_Information

## Acknowledgements

This work was funded by the Novo Nordisk Foundation Copenhagen Bioscience Ph.D. Program grant No. NNF22SA0078231 (awarded to M.H.) and Novo Nordisk Foundation grant No. NNF20CC0035580. N.G.M. was supported by the Danish Data Science Academy, which is funded by the Novo Nordisk Foundation (grant No. NNF21SA0069429) and VILLUM FONDEN (grant No. 40516). A.C.A.A. was funded by the Rubicon research programme which is financed by the Dutch research council (NWO) (file number 019.231EN.007). J.A.A. received funding from the Novo Nordisk Foundation grant No. NNF24OC0095360.

