## Supplementary material for "Biocatalytic Production of Galantamine in Yeast through Cytochrome P450 Optimization": Holtz et al, 2026_Supplementary_Information

### Contents of Supplementary Information

**Supplementary Figure 1.** LC-MS/MS detection of putative galantamine pathway intermediates produced in yeast.

**Supplementary Figure 2.** Extracellular pH affects the intracellular detection of 4OMe-norbelladine by a RamR-based biosensor.

**Supplementary Figure 3.** Computational attribution of galantamine MS/MS fragments using SIRIUS.

**Supplementary Figure 4.** Accumulation of galantamine pathway intermediates in engineered yeast strains.

**Supplementary Figure 5.** LC-MS/MS characterization of putative reduced shunt products of the galantamine pathway in yeast.

**Supplementary Figure 6.** Detection of putative 8-fluoronarwedine following bioconversion of 6′-fluoro-4OMe-norbelladine in engineered yeast.

**Supplementary Table 1.** Codon optimized gene sequences cloned in this study.

**Supplementary Table 2.** Plasmids used in this study.

**Supplementary Table 3.** Strains used in this study.

### Supplementary Figures

A)

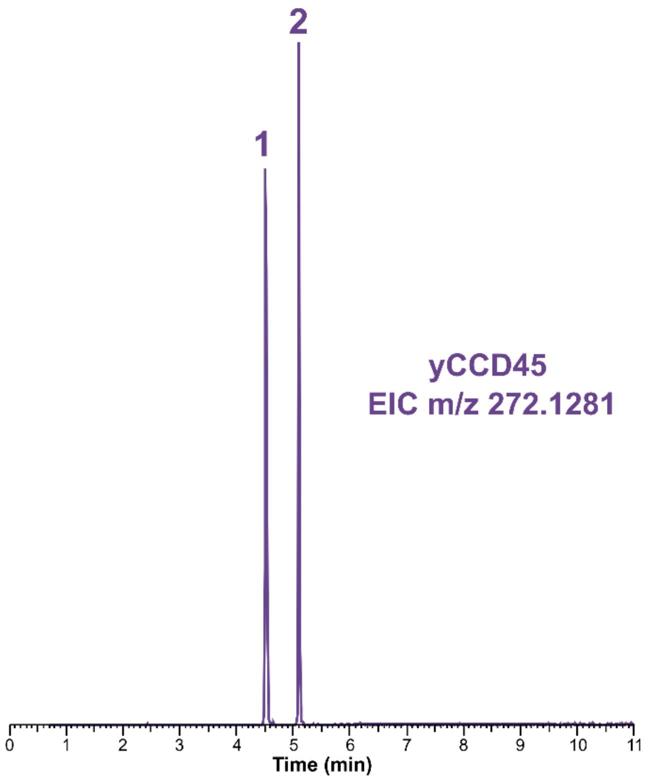


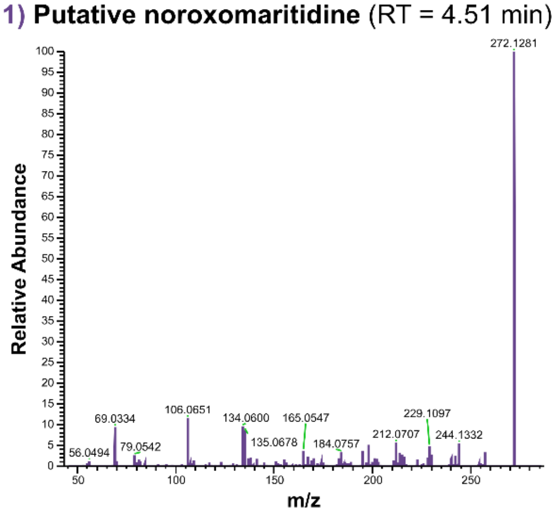


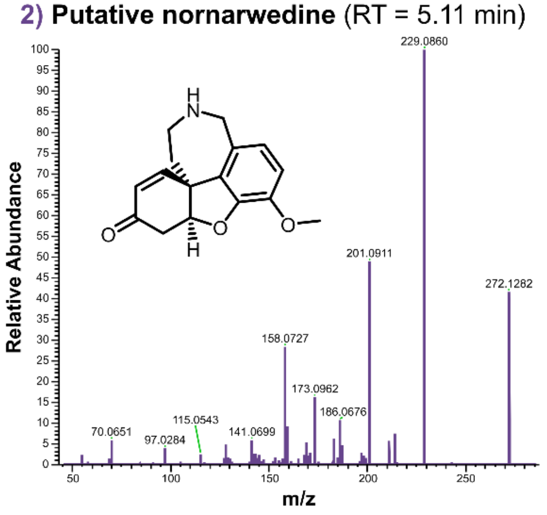


B)

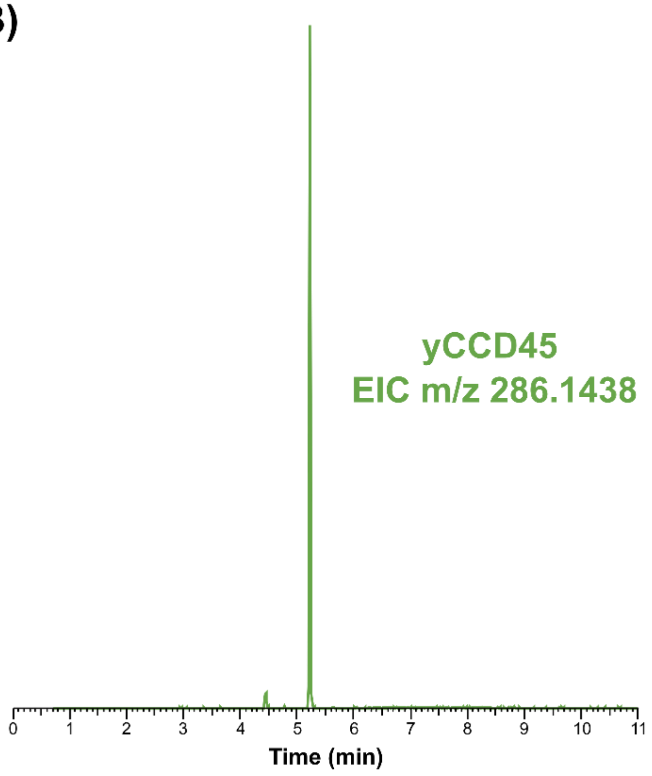


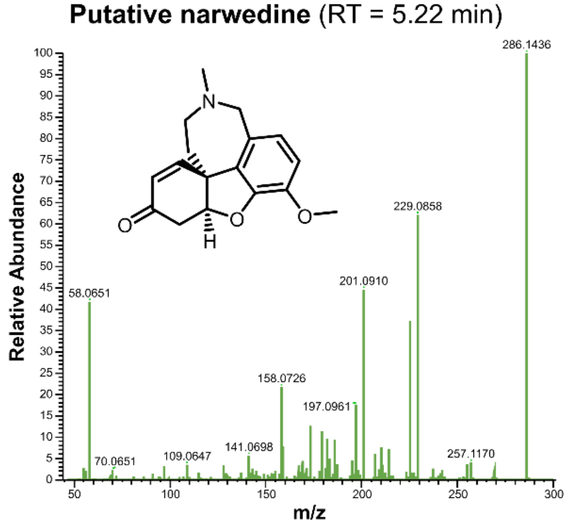

C)

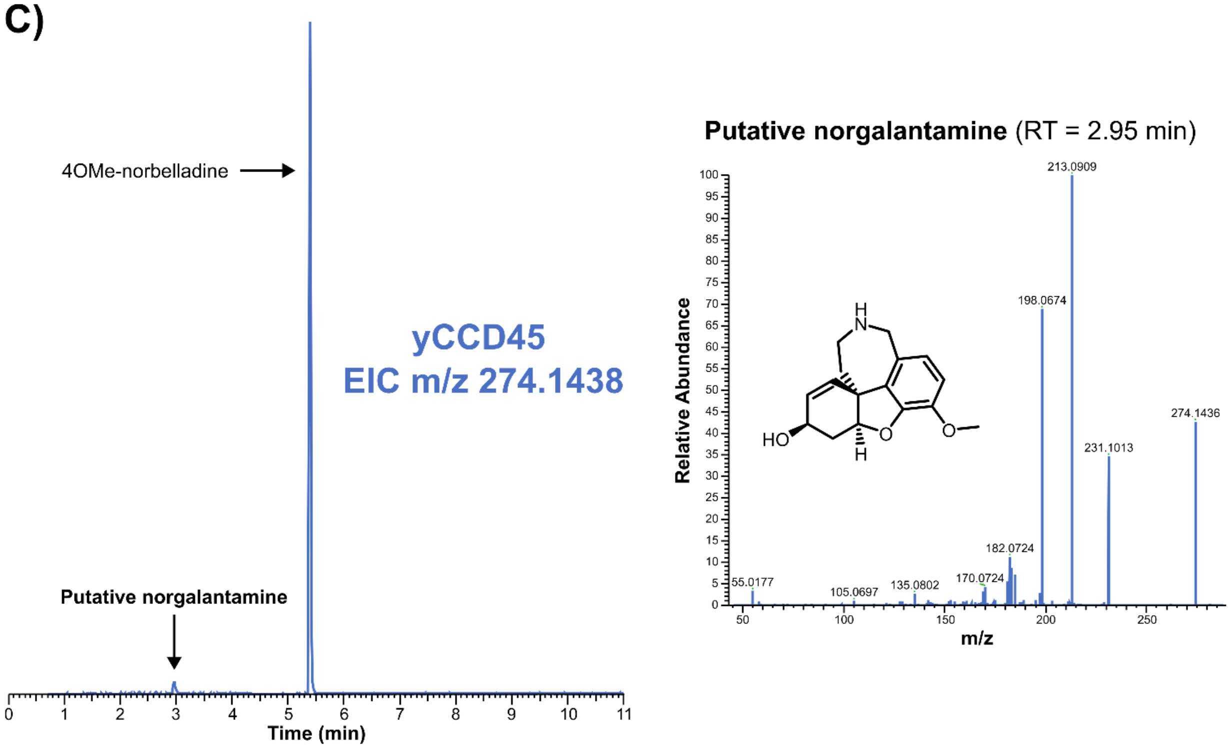


**Supplementary Figure 1. LC-MS/MS detection of putative galantamine pathway intermediates produced in yeast.** Engineered yeast strain yCCD45 was cultivated in synthetic complete medium containing 2% (w/v) glucose at 30 °C for 6 days. Extracted-ion chromatograms and corresponding MS/MS spectra are shown for putative pathway intermediates **A)** nornarwedine m/z 272.1281, **B)** narwedine m/z 286.1438 and **C)** norgalantamine m/z 274.1438 (5 ppm mass resolution). No GAL was detected in these conditions. Metabolite attribution was supported by SIRIUS analysis of the MS/MS fragmentation spectra.

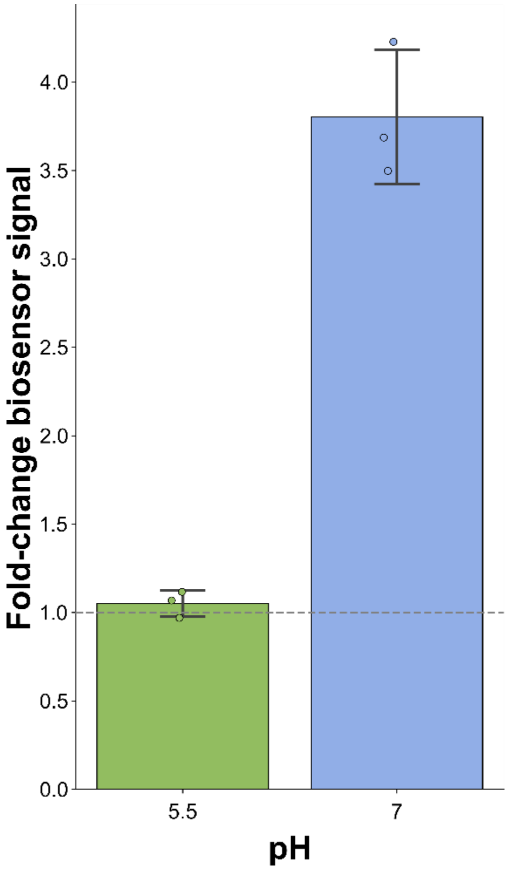


**Supplementary Figure 2. Extracellular pH affects the intracellular detection of 4OMe-norbelladine by a RamR-based biosensor.** A yeast strain carrying the intracellular 4OMe-norbelladine biosensor was supplemented with 250 µM ligand at the indicated pH values in synthetic complete media with 2 % glucose. Biosensor output was quantified as the median fluorescence intensity (mMFI) of the cell population measured in the FITC-H channel using flow cytometry, corresponding to yeGFP fluorescence. The signal was normalized to the corresponding condition without ligand and is reported as fold change. The greater response observed at pH 7 is consistent with enhanced uptake and intracellular availability of the pathway substrate 4OMe-norbelladine. The dashed horizontal line denotes a fold change of 1. Individual data points represent independent biological replicates, bars and error bars show the mean ± SD (n = 3).

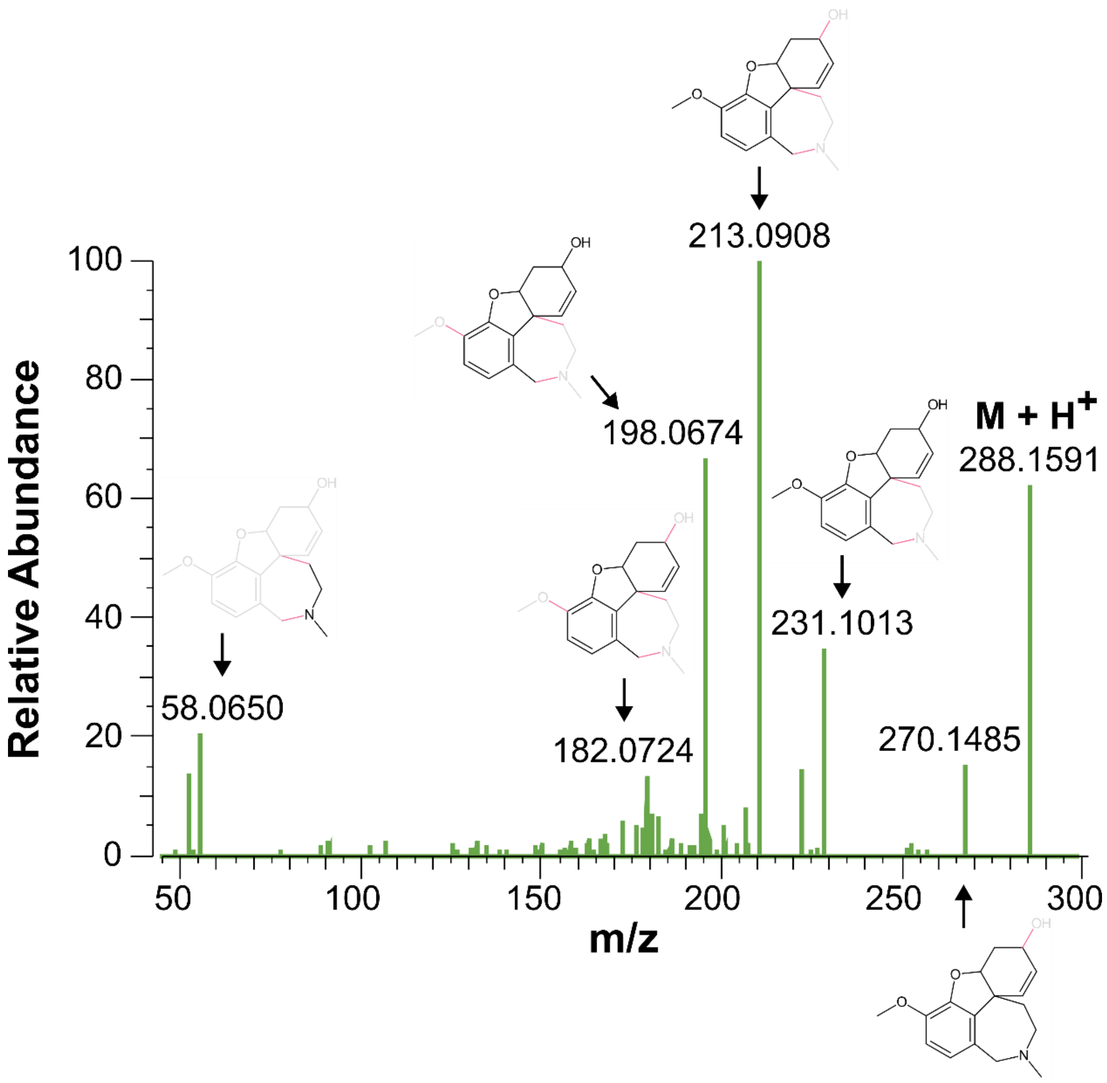


**Supplementary Figure 3. Computational attribution of galantamine MS/MS fragments using SIRIUS**. The experimental MS/MS product-ion spectrum of an authentic galantamine standard is shown together with the proposed structures of selected fragment ions. Red bonds within the molecular structures indicate the bond cleavages associated with the proposed fragmentation pathways.

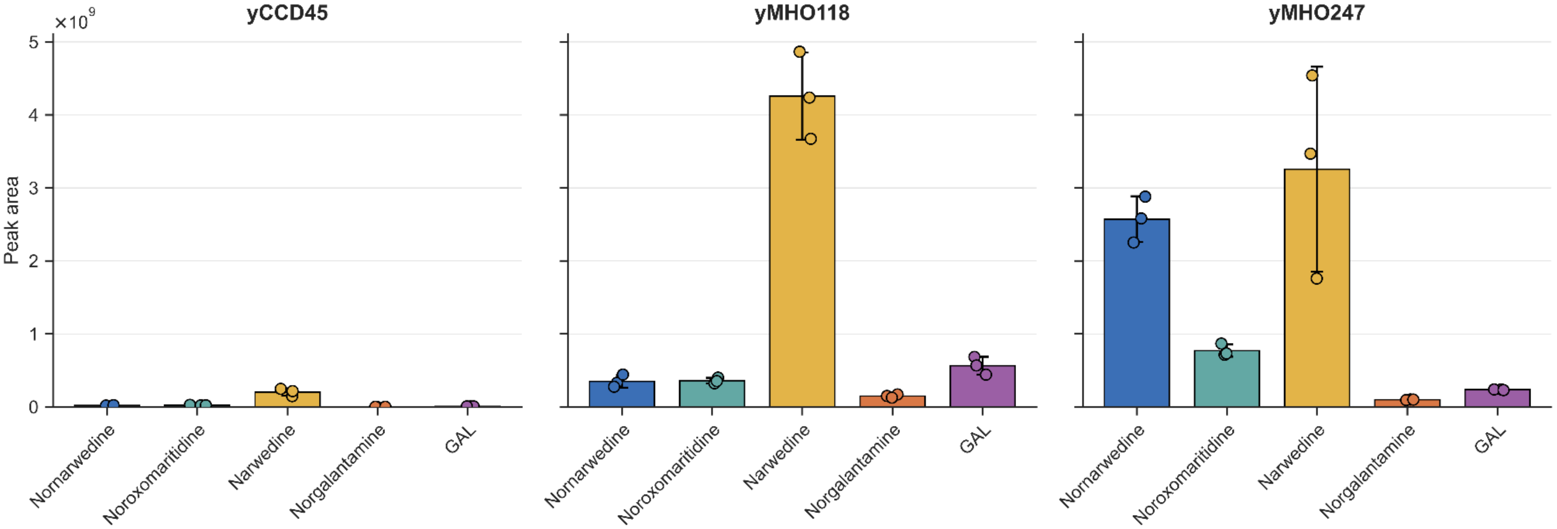


**Supplementary Figure 4. Accumulation of galantamine pathway intermediates in engineered yeast strains.** Strains yCCD45, yMHO118, and yMHO247 were cultivated in synthetic complete medium containing 2% (w/v) trehalose at pH 7 and 25°C for 6 days. yCCD45 lacks heterologous cytochrome P450 reductase (CPR) and cytochrome $b_5$ (CYB5), yMHO118 expresses NpCPR and CrCYB5, and yMHO247 was derived from yMHO118 by introducing a high-copy 2 μ plasmid overexpressing NtNMT1 and NtAKR1. Individual data points represent independent biological replicates, bars and error bars show the mean ± SD (n = 3).

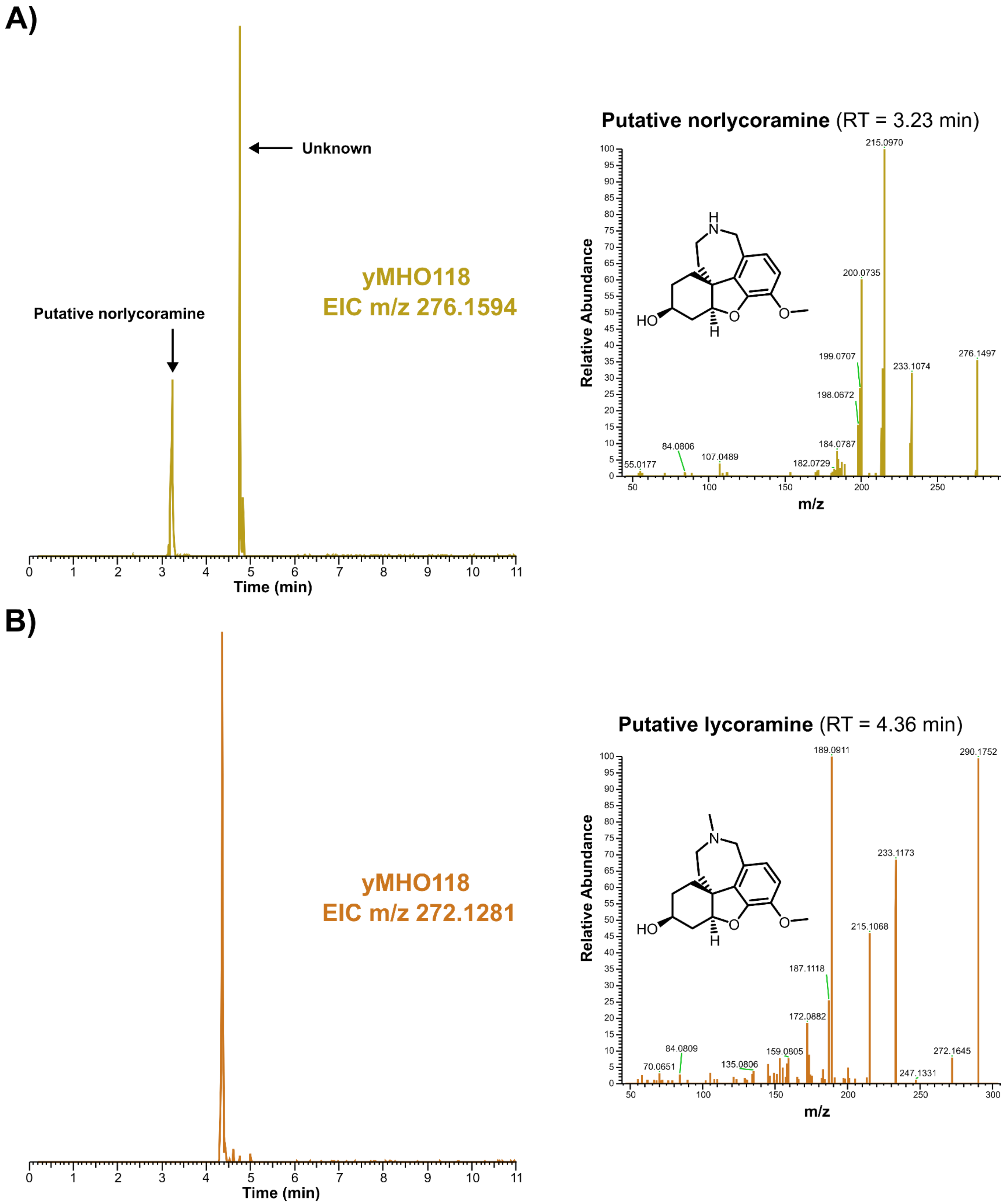
A)
Unknown
Putative norlycoramine
yMHO118
EIC m/z 276.1594
Time (min)
Putative norlycoramine (RT = 3.23 min)
Relative Abundance
m/z
215.0970
200.0735
199.0707
198.0672
184.0787
182.0729
233.1074
276.1497
107.0489
84.0806
55.0177
HO
O
O
H
N
H
B)
yMHO118
EIC m/z 272.1281
Time (min)
Putative lycoramine (RT = 4.36 min)
Relative Abundance
m/z
189.0911
290.1752
233.1173
215.1068
187.1118
172.0882
159.0805
135.0806
84.0809
70.0651
247.1331
272.1645
HO
O
O
H
N

**Supplementary Figure 5. LC-MS/MS characterization of putative reduced shunt products of the galantamine pathway in yeast.** Extracted-ion chromatograms and corresponding MS/MS product-ion spectra are shown for metabolites detected at **A)** m/z 276.1594 and **B)** m/z 272.1281 in strain yMHO118. Based on accurate mass and MS/MS fragmentation, these metabolites were putatively attributed to reduced forms of norlycoramine and lycoramine respectively.

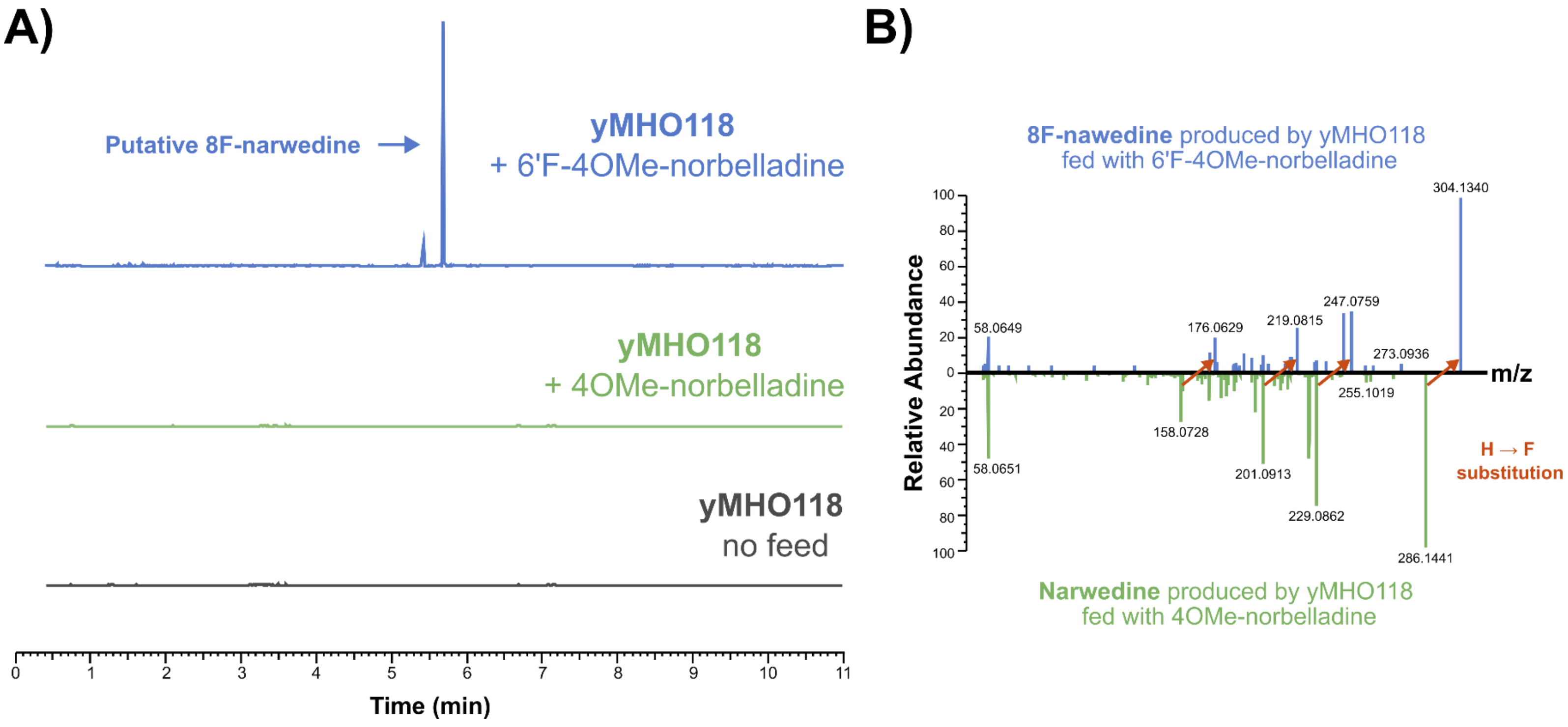


**Supplementary Figure 6. Detection of putative 8-fluoronarwedine following bioconversion of 6′-fluoro-4OMe-norbelladine in engineered yeast. A)** Extracted-ion chromatograms (m/z 304.1343) showing a feature assigned as putative 8-fluoronarwedine in strain yMHO118 supplemented with 6′-fluoro-4OMe-norbelladine (blue). The corresponding feature was not detected in yMHO118 supplemented with nonfluorinated 4OMe-norbelladine (green) or in the no-feed control (gray). **B)** Mirror MS/MS comparison of putative 8-fluoronarwedine (blue) and narwedine (green). Orange arrows connect corresponding product ions exhibiting the expected +17.9906 m/z shift associated with H-to-F substitution in fluorine-retaining fragments.

### Supplementary Tables

**Supplementary table 1. Codon optimized gene sequences cloned in this study** (start codons are highlighted in bold; stop codons are underlined).

| Name | Organism | Reference | Nucleotide sequence |
|---|---|---|---|
| NtCYP96T6 | *Narcissus cv. Tête-à-Tête* | [1] | **ATG**GTCACCTCTTCCTCTGCTTGGTTAATCTTCTCTGACCACTACAGAGAAATCTTG<br>ATCGCCATTGCTTGTTTGGTTGTGTTCTCCCTATTAAGATCTGCCAGATCTTCTTCC<br>AAGGGCGGTCTACCATACAACTGTCCAATCTTCGGTATGTTGCCAGCTATCATCTC<br>CAACAACCAATTCAACGATTTTATCACCGCTCGTTTGAGAGAAATCGGTTGGACCT<br>TCAGCTTCAAGGGTCCATGGTTGTTGGACATGGACTACATCTTTACTTGTGATCCA<br>GCTAACATCAACCACATGTTCAACCACAACTTTGAAAACTATCCAAAGGGTCAATTA<br>GGTGAAGTTTTCGATATTTTCGGTAACAACATTTTGAACGCCGATGGTGACGTCTG<br>GAGAAACCATAGAAAGATGGCTCAAACTATCATGTGGGACGAAAACTACAGAACCA<br>TGCAAGCCACTTTCATCAGAAACAAGATGGAAGACGCTTTAATCCCAGTCTTAGAT<br>TCAGCTGCTTGTGAACGTAAGCCAGTTGACTTGCAAGATGTTTTCTTGAGATTTACT<br>TTTGACACAAGTTGTTTCTCTGTTTTGGCCGCTGACCCAGAATCTTTGACTATGGAA<br>TTCCCAACCGTTCCTTTCCCACAAGCTGTCGACCAAGCTTTGGATGCTGCTTTGAG<br>AAGACACATTATGCCTAGATTGATTTGGAAGTTGAAAAGATTCTTCAAGATTGGTTC<br>CGAACGTACTTTGGCTACTGCTTGCGAAGTCATTGACTCCTACTTGTACGAAAAGA<br>TCGCTCAATTGAAGGCTAACAGAGTTGGTATGGGTAAGATCAAGTCTTACGATGTT<br>GTTTCCTTCTACATGGATAACTTCGACATTCACGACGACAAGTTCTTGAGAGACAAT<br>GCCTTCACTTACCTGCTGTTGCAAAGAAACACTCAATCAATTACCATGACCTGGTT<br>GTTCTACGCTTTGTTCGAAAATCCAAAGGTTGAACCTAAAATTTTGATTGAATTAAA<br>GTCTATCGTCGATGAATCCTCCGAGCGGAAATTGAAGGACGGTTTCACTTTGTTTG<br>ATTCTAACATGATCCAATCCGCTATTTACTTGCACGCTACCTTGTGTGAAGCTTTGA<br>GAATCTACCCACCAGTTCCATTCGAAATTAAGAACGCTCACGAAGCTGATGTCTTG<br>CCATCTGGTCACAAGGTCAGAGCTGGTGAAAAAATTTTATACTCTCCATATGCTATG<br>GCCAGAATGACTGGTATATGGGGTGACGATTGTTTGGAATTCAAGCCAGAAAGATG<br>GATCACTGACAACGGTATGCTAAAGCACGAACCAGCTTACAAGTTTTTCTCTTTCAA<br>TGCCGGTCCAAGAATCTGTGTCGGTAAGGAACTCTCTTTCACCCAAATGAAAATGG<br>TTGCTGCCACCATTATTTACAACTTCCATGTTCAAATGGTCAAGGGACATGTCGTC<br>GAACAATCTAACTCCATTTTGATGGAAATGAAGCACGGTTTGATGGGTGAAGTTATT<br>AAGAGATCTGTTATG<u>TGA</u> |
| NtNMT1 | *Narcissus cv. Tête-à-Tête* | [1] | **ATG**GACGAAAGAACCAAGGGTATGGTTGCTTTCTTCGACGAAACTGCTATGCACGA<br>CTCTTTGTGGATGGAAGACTTGCATCACGGTTACTACGGTCCAGGTGAACAAGCTC<br>ACGTTTCTACTAACCATGCCGCTGAAAAGAGAATGGTCGAAGAAGTTTTGAAGTTT<br>GCTGGTGTCTCCGATGACCCAATCAAGAAGCCAAAGAACATCATTGATGTTGGTTG<br>TGGTTTCGGTGGTGCCGCTATTCATTTGTCCAAGAAGTACGGTGCTAACTGTACTG<br>GTATCAACTTGTCTCCAGCTCAAATCCAAAAGGCTAAGAATTTAGCTGACGCCAAC<br>GGTTTGGGTGACAAGACCTCTTTCGTTGTCGCTGATGCTTTAAACCAACCATTCCC<br>AGATGGTCAATTCGATTTAGTCTGGTCTATGGAAGTCATTGAACACACTCCAGACA<br>AGTTGAAATTCATCTCTGAAATGGCTAGAGTTGCCGCTCCAGGTGCTACCATCATC<br>TGCACCTTGTGTTGTCACAGAGACTTGTCCCCAGGTGAAAAGAGTTTGAAGCCTGA<br>TGAAAAGAACTTGATTGACAAAATTTTGAAGACTTTCCACCAACACTCTTACATTTCT<br>CCATCTGACACAGTCAAGATCGTTCAATCCTTGGCTTTGCAAGATATCAAGATTGC<br>CGATTTTTCCGAAAACGTCTTGCCATACATCAGAGCTAATGTCCAATCTCAAAGAAC<br>CTGGAAGGGTCTAGCCTCGATGGTTTTGAACGGCTGGACCAACTTCAAGGTTTCC<br>CAAAACCAACCTTTGACTCTTAAAACTTACGAAAAAAACTTGACCCGTTATACTGTT<br>ATTGCTTGTCACAAGCCAAAG<u>TGA</u> |
| NtAKR1 | *Narcissus cv. Tête-à-Tête* | [1] | **ATG**TTGAACAACATTCCAGAATTGATTCTTACCAAGGATTCTAGAGCTATGCCAGTC<br>GTCGGTATGGGTACCGCTACTCACCCTTTCCCACAAGATAACACTAAATCTGCTAT<br>CATGGATGCTATTGAAATCGGTTACAGACACTTCGACACTGCCTCTTTGTACGGTT<br>CTGAAGAACCATTGGGTGAAGCAATGGTCGAAGCCCAAAAGTTGGGTTTAATCAAA<br>TCCAGAGAAGAATTATTCATCACTTCCAAGTTGTGGTGTAATGAAGCTCACCCAGG<br>TTTGGTTATCCCAGCCATCAAGAAGAGTCTATTGAATTTGAAGTTGGACTACTTGGA<br>TTTGTATCTCATTCACATGCCATTCACCACCAAGCCAGAATCTCCACCATTCCCATT |

| | | | |
|---|---|---|---|
| | | | GTACAAGGAAGACATGGTTACCATGGACTTGCAAGGTGTCTGGAAGGCTATGGAA<br>GAATGTAAGAGATTGGGCATGGCTAAGGCCATTGGTGTTTCCAACTTCACTGTTAA<br>CAAATTAGAAGAATTGTTGCCATTTGCTAAGATTCCTCCACAAGTCAACCAAGTTGA<br>AATGAACCCAGCTTGGCAACAACAAAAGTTAAGACAATACTGTAACGCCAAGGGTA<br>TCCATGTTACTGCTTACTCCCCCTTAGGTGGTCAAGAGGAACCATCTTCTCCAAAC<br>ATGGTTATGAAGTCAGAAGATTTGAAGCAAATTGCTAACGCTAGAGGTAAGACCGT<br>TGCTCAAGTTTCTTTGAGATGGGTTTACGAACAAGGTGTCTCCATCGTTGTCAAGT<br>CTTTCAACAAGGAACGTATCAAGAAGAACATCGAAATTTTTGACTGGAACTTGTCC<br>GAAGAAGAATGTCACATGATTTCTCAAATCCCACAATGCAAAAAGACTACAGTCGA<br>AGCTATGTTCTCCAACTGGAAGCCACATCCAGACGAATACTTCTTCGAAGAAAACG<br>AAGGTGAATGA |
| RamR 4NB2 | *Salmonella typhimurium* | [2] | **ATG**GTTGCTCGCCCAAAGTCTGAGGACAAAAAGCAGGCATTGCTTGAAGCGGCAA<br>CTCAAGCCATCGCGCAATCAGGCATTGCCGCTAGTACCGCTGTAATTGCACGCAA<br>TGCGGGAGTTGCGGAAGGGACGTTGTTCCGCTATTTCGCAACGAAAGATGAGTTG<br>ATCAACACCCTTTACTTACATTTGACTCAGGACATGTGCCAATCAATGATCATGGAA<br>TTGGATCGTTCTATTACTGACGCTAAGATGATGACCCGTTTTATCTGGAACAGTTAT<br>ATTAGCTGGGGATTGAACCACCCAGCTCGCCATCGTGCCATTCGTCAGTTGGCGG<br>TTTCTGAAAAGTTGACGAAGGAAACCGAACAACGCGCGGATGATATGTTCCCGGA<br>GTTACGCGACTTGGATCACCGTGGTGTTCTTATGGTGTTTATGTCCGACGAGTACC<br>GCGCCTTCGGCGACGGGTTGTTCTTGGCGCTTGCTGAGACGACTATGGATTTCGC<br>TGCGCGCGACCCGGCTCGCGCTGGTGAGTACATTGCGTTGGGCTTCGAGGCTAT<br>GTGGCGCGCACTTACGCGCGAAGAGCAGTAA |
| AtATR1 | *Arabidopsis thaliana* | [3] | **ATG**ACATCAGCTCTTTACGCCTCTGATCTATTTAAACAGCTGAAATCTATTATGGGA<br>ACAGACTCATTGTCAGATGATGTTGTTTTAGTGATCGCTACCACCTCATTGGCTCTG<br>GTAGCCGGCTTTGTCGTACTATTGTGGAAAAAAACTACCGCAGATAGATCTGGTGA<br>GTTGAAACCCTTGATGATTCCGAAATCACTGATGGCTAAAGACGAAGATGATGACT<br>TAGACTTGGGTTCAGGTAAAACGAGAGTGTCTATCTTCTTCGGTACTCAGACCGGA<br>ACAGCTGAAGGTTTTGCCAAAGCACTTTCTGAAGAAATTAAAGCAAGATATGAAAAA<br>GCTGCTGTTAAAGTGATCGATTTGGATGATTATGCCGCTGACGACGATCAATACGA<br>AGAAAAGTTAAAAAAGGAGACATTAGCATTCTTCTGCGTGGCTACTTACGGTGATG<br>GTGAACCTACTGATAACGCCGCGAGATTTTACAAGTGGTTCACAGAAGAAAACGAA<br>CGTGACATCAAATTGCAGCAATTGGCATATGGCGTCTTTGCACTGGGAAACAGACA<br>ATATGAACACTTCAACAAAATTGGTATTGTGTTGGATGAAGAACTTTGTAAAAAAGG<br>AGCCAAGAGGTTGATTGAAGTTGGTCTAGGCGACGATGATCAATCAATAGAGGAT<br>GACTTTAACGCATGGAAAGAGTCCTTGTGGTCTGAATTGGACAAACTTTTGAAAGA<br>CGAGGACGATAAGTCTGTTGCGACCCCTTATACGGCTGTGATACCTGAATATAGAG<br>TTGTGACGCATGATCCTCGTTTTACGACACAGAAGTCAATGGAGTCCAACGTTGCG<br>AATGGAAACACCACAATTGACATTCACCATCCCTGTAGAGTTGATGTTGCCGTTCA<br>AAAGGAGCTTCATACGCACGAAAGTGACAGATCTTGCATACACTTAGAATTTGATAT<br>TTCACGTACTGGTATTACTTACGAAACCGGAGATCACGTGGGTGTCTATGCTGAAA<br>ACCATGTAGAGATCGTCGAGGAGGCAGGTAAATTGTTAGGGCACAGTCTTGATTTG<br>GTATTTTCTATCCATGCCGATAAAGAAGATGGTTCACCCTTAGAATCCGCAGTGCC<br>ACCGCCTTTCCCAGGTCCATGCACTTTGGGAACGGGTTTGGCTAGGTACGCTGAC<br>TTGTTGAATCCACCAAGGAAATCAGCTTTGGTGGCTCTTGCAGCGTACGCAACCGA<br>GCCATCAGAAGCAGAAAAATTAAAACATCTTACCTCTCCAGATGGAAAAGATGAAT<br>ACTCTCAGTGGATAGTGGCATCCCAACGTTCATTGTTAGAAGTAATGGCGGCCTTC<br>CCATCAGCTAAACCGCCATTGGGTGTTTTTTTCGCCGCCATAGCCCCGAGGTTACA<br>ACCTAGGTACTATTCCATTTCTTCATCTCCAAGGTTAGCTCCCAGTAGAGTGCACG<br>TTACTTCAGCTCTTGTATATGGTCCAACACCTACTGGTAGAATTCATAAGGGGGTG<br>TGTAGCACATGGATGAAAAACGCTGTACCAGCAGAAAAATCACATGAATGTTCTGG<br>TGCGCCAATTTTTATAAGAGCTTCAAACTTTAAATTGCCTTCTAACCCATCAACACC<br>TATTGTAATGGTTGGACCTGGTACAGGATTGGCACCATTTAGGGGTTTTTTGCAAG<br>AAAGAATGGCCTTGAAGGAAGACGGGGAAGAATTAGGTTCTAGCTTGTTGTTCTTC<br>GGATGTAGAAATAGACAGATGGATTTCATCTACGAAGACGAATTAAACAATTTTGTA<br>GATCAAGGTGTTATTTCTGAACTTATCATGGCGTTTTCTAGAGAGGGCGCTCAAAA<br>AGAGTACGTACAGCATAAGATGATGGAAAAGGCTGCTCAAGTTTGGGATTTAATTA<br>AAGAAGAAGGCTATCTATACGTCTGTGGAGACGCAAAGGGTATGGCGCGTGATGT<br>TCATAGGACATTACATACTATAGTCCAAGAACAAGAAGGTGTCAGTTCTTCTGAAG |

| | | | |
|---|---|---|---|
| | | | CCGAGGCCATTGTTAAAAAGTTACAAACTGAAGGCAGATACCTTAGAGATGTCTGG<br>TAA |
| CrCYB5 | *Catharanthus roseus* | [3] | **ATG**GCCTCTGATCAAAAGTTGCATAAGTTCGATGAAGTCTCAAAACATAATAAAACG<br>AAAGATTGTTGGCTGATTATTAATGGTAAGGTCTACGACGTCACTCCGTTTATGGA<br>CGATCATCCAGGTGGTGACGAAGTCTTATTATCCGCCACAGGCAAGGACGCAACA<br>AATGACTTTGAAGATGTTGGTCACTCTGACAGCGCTAGAGAAATGATGGATAAATA<br>TTACATTGGTGAGATGGATATGGCTACTGTTCCACTTAAAAGAACATACATTCCTCC<br>ACAGCAAGCTCAATATAATCCTGACAAGACACCAGAGTTCGTGATTAAGATCCTTC<br>AATTTTTAGTACCCTTGCTGATATTGGGTTTAGCGTTCGCTGTTAGACATTACACCA<br>AGGAAAAATAA |
| LrCPR2 | *Lycoris radiata* | [4] | **ATG**AAACCATCACCATTGTCCTTGTTGATGGCTATCAGTACTGGTAACTTGGGTGA<br>TGGTGGTTTGCCACCAGAAGTTGCTGCCATCGCCGAAAACAGAGAAGTTTTGATGA<br>TCTTGACCACCTCCGTTGCTGTCTTCGTCGGTTGTGTTGCTTTGTTCTTGTGGAGA<br>AGATCTTCTGCCAAGTCTTCTAAGTCTTTGGAACCTTTGAAGCCAATGGTTGTCTCC<br>AAGGAACCAGAAATTGAAGTCGACGATGGTAAGAGAAAGGTTACTATTTTCTTCGG<br>CACTCAAACTGGTACCGCTGAAGGTTTCGCTAAAGCTTTGGCTGAAAAGGCTAAGG<br>CTGGTTACGACAAGGCTGCTTTCAGAATTGTTGATTTGGACGACTACGCTGCTGAT<br>GATGATGAATACGAGGAAAAATTGAAGAAGGAAACTTTGGCTTTATTCTTTTTGGCC<br>ACCTACGGTGACGGTGAACCAACTGACAATGCTGCTAGATTCTACAAATGGTTCAC<br>CGAAGGTAAGGAAAGAGCTAAGTGGTTGGAAAATCTTCAATACGCCGTCTTCGGTT<br>TGGGTAACAGACAATACGAACACTTCAACAAGGTCGGTAAGGTTGTTGACGAAATC<br>TTAGCTGAACAAGGTGCTAAGAGATTGGTTCCAGTTGGTTTAGGTGATGATGACCA<br>ATGTATCGAAGATGATTTCACTGCTTGGTCTGAACTCTTATGGCCAGAATTAAACAA<br>GTTGTTGAGAGATGAAGATGACTCCTCCGGTGCTATCACCACTTACACTGCCGCCA<br>TCCCAGAATACAGAGTTGTTTTCATTGAACCAGGTACCTCTCCATTGGAAAAGAACT<br>GGTCTTTGGCTAACGGTCATGCTGTTCACGACATCCACCACCCATGTCGTGCCAAC<br>GTCGCCGTCAGACGTGAATTGCACACTCCAGCTTCTGACAGATCTTGTGTCCATTT<br>GGAATTCGACATTGCTGGTACTGGTTTGGTCTATGAAACCGGTGACCACGTTGGTG<br>TCTACTCCGAAAACTGTTTGGAAACCGTTGAAGAAGCTGAAAAAATGTTGGGCTTA<br>TCTCCAGACACTTTCTTCTCCATTCACGCTGACAACGAAGACGGTACTCCATTGTC<br>TGGCACTTCTCTTCCACCTCCATTCCCATCGCCATGCACTTTGAGAACTGCTTTGA<br>CCAGATATGCTGATTTGTTGAACAGCCCAAAGAAAGCAGCTTTGATTGCTTTAGCT<br>GCCCATGCCTCCGACCCAAACGAAGCTGAAAGATTGAGACACTTGGCTTCTCCAG<br>CTGGGAAGGACGAATACAGTAGATGGATTGTTGCCTCTCAAAGATCCTTACTAGAA<br>GTCATTGCTGAATTCCCATCTGCCAAGCCACCATTAGGTGTTTTCTTTGCTGCTATT<br>GCTCCAAGATTGCAACCAAGATACTACTCTATTTCATCCTCTCCAAGAATGGCTCCT<br>ACTCGTATCCATGTCACGTGTGCTTTGGTATACGCTCCAACCCCGACCGGTAGAAT<br>TCACAAGGGTGTCTGTTCCACCTGGATGAAGCACGCCGTTTTGATGGAAGAATCTG<br>AAGAATGTTCTTGGGCTCCAATCTTTGTTCGTCAATCTAACTTCAAGTTGCCATCCA<br>ACCCATCTACACCAATCATCATGATTGGTCCAGGTACCGGTTTGGCTCCATTCAGA<br>GGTTTCTTGCAAGAAAGATTAGCTTTGAAGGAAGCAGGTACTGAACTGGGTCCAGC<br>CATTTTCTTTTTTGGTTGCAGAAACAGAAAAATGGACTTCATCTACGAAGAAGAATT<br>GTACAACTTTGCCGAAGCCGGTGCTGTTTCCGAGTTAATCGTCGCATTCTCTAGAG<br>AAGGTCCAACCAAGGAATATGTTCAACACAAGATGGCTGAAAAGGCTCCGGAATTG<br>TGGAACATCATCTCCAATGGTGGTTACATTTACGTTTGTGGTGATGCTAAGGGTAT<br>GGCCAGAGATGTTCACAGAGTTTTCCACACCATCGTCCAAGAGCAAGGTTCTTTGG<br>ATGGTTCCAAGGCTGAATCCATGGTCAAGTCTTTGCAAATGGAAGGTCGTTACTTG<br>CGTGATGTCTGGTAA |
| NpCPR | *Narcissus pseudonarcissus* | This study. | **ATG**CACCCAGAAACCATGAAACCATCTTCTTTGTCTTTGTTGACTGCTATTTTCACC<br>GGTAACTTGGGTGATGGCGGGTTGCCTCCAGAAATGGAAGCTATCGCCAAGAACA<br>GAGATGTTTTGATGTTGTTGACCACTTCTATTGCCGTCATCGTTGGTTGTGTCGTCT<br>TCTTCCTCTGGAAGAGATCTTCGTCTAAAAGTTCTAAGTCCTTTGAACCATTGAAAC<br>CAATGATGGTTTCCAAGAAGCCAGAATTTGAAGTTGATGATGGTAAGAAGAAGGTT<br>ACCGTTTTCTTCGGTACTCAAACTGGTACCGCCGAGGGTTTCGCTAAGGCTCTAGC<br>TGAAGAAGCCAAGGCCAGATATAATAAGGCTTCCTTCCGTATCGTCGATTTGGACG<br>ACTACGCTGACGATGATGATGTCTACGAAGAAAAGATGAAGAAGGAAACCTTGGCT<br>TTATTCTTTTTGGCCACTTACGGTGACGGTGAACCAACTGACAACGCTGCTAGATT<br>CTACAAGTGGTTCACCGAAGGTAAGGAAAGAGAAAAATGGTTGGAAAACTTGCAAT<br>ACGCCGTTTTCGGTTTGGGAAACCGTCAATATGAACACTTCAACAAGGTTGCTAAG |

| | | | |
|---|---|---|---|
| | | | GTTGTTGACGAAATCTTGGCTGAACAAGGTGCCAAGAGATTGGTTCCAGTCGGTTT<br>GGGTGACGACGACCAATGTATTGAAGATGATTTCACTGCTTGGAGAGAATTGTTGT<br>GGCCAGAATTGGACAATTTATTACGTGACGATGACTCCCTAGGTGCTACCACCACC<br>TACACCGCTGCCATTCCAGAATACAGACTGGTCTTGATTGATTCTGACGCTTCTCC<br>TTTGGAGAGAAACTGGTCTTTAGCTAACGGTCACGCCGTGCACGACATCCACCAC<br>CCATGTAGAGCTAACGTTGCTGTCAGAAGAGAATTGCACACTCCAGCTTCTGACAG<br>ATCTTGTATCCATCTAGAATTCGACATCGCTGGTTCTGGTTTAGTCTACGAAACTGG<br>TGACCACGTCGGTGTTTACTCTGAAAACTGTTTGGAAACAGTTGAAGAAGCTGAAA<br>AGTTGCTAGGTCTATCCCCAGACACTTTCTTCTCCATTCATGCTGATAACGAAGATG<br>GTACTCCATTGTCCTGCACTTCCTTGCCACCACCTTTCCCATCTCCATGCACTTTGA<br>GAACTGCTTTAACAAGATACGCTGACTTGTTGAACTCTCCAAAGAAGTCTGCTTTGT<br>CTGCTTTGGCCGCTCACGCAAGCGACCCAAACGAAGCAGAAAGATTGAGACACTT<br>AGCTTCTCCAGCTGGCAAGGATGAATACTCTCAATGGATTGTCGCTTCCCAAAGGA<br>ACTTGTTGGAAGTCATGGCCGAGTTTCCAAGTGCTAAGCCACCTTTGGGTGTCTTT<br>TTTGCGGCTATCGCTCCACGTTTGCAACCAAGATACTACTCCATCTCCTCCTCTCC<br>GAGAATGGCTCCAACCAGAATCCATGTTACTTGTGCTTTGGTCTATGGTCCAACCC<br>CAACTGGTAGAATTCACAAGGGTGTTTGTTCTACCTGGATGAAGCACGCTGTTCCA<br>ATGGAAGAATCTGAAGACTGTTCTTGGTCCCCAATCTTCGTTAGACAATCAAATTTC<br>AAATTACCATCCAACCCATCCACTCCAATTATCATGATTGGTCCAGGTACTGGTTTG<br>GCACCATTCAGAGGTTTCTTGCAAGAAAGATTGGCTTTGAAGGAAGCCGGTACTGA<br>ATTGGGTCCAGCCATTTTGTTCTTCGGTTGTCGTAACAGAAAGATGGATTTCATCTA<br>CGAAGAAGAATTGTACAACTTTGCTGAAGCTGGTGCTGTTTCTGAATTAATTGTTGC<br>TTTCTCTAGAGAAGGTCCAGCTAAAGAATACGTTCAACACAAGATGACCGAAAAGG<br>CTCCTGAATTGTGGAACATTATCTCTAACGGTGGTTACATCTACGTCTGTGGTGAT<br>GCTAAGGGTATGGCCCGTGATGTTCACAGAGTTTTGCATACCATCGTCCAAGAACA<br>AGGTTCCTTGGACTCGTCTAAGGCTGAATCCATGGTCAAGTCCTTACAAATGGAAG<br>GTCGTTACTTGAGAGATGTTTGGTGA |
| NpCYB5 | *Narcissus pseudonarcissus* | This study. | **ATG**GCTGCTGAATCTAAGGTCTACCACTTCGAAGAAGTTTCCAAGCACAACGCCAC<br>CAAGGACTGTTGGTTAATCATCAACGGTAAAGTCTATGATGTCACTCCATTCATGG<br>AAGACCATCCAGGTGGTGACGAAGTCTTGTTGGCCGCCACTGGTAAGGATGCTAC<br>CAATGATTTTGAAGATGTTGGTCACTCAGATGGTGCTCGTGACATGATGGGTAAGT<br>ACTTCATCGGTGAAATTGACGCTTCCACTGTTCCAACTAAGAGATCTTACGTTGCTC<br>CACAACAACCAGCTTACAACCCAGACAAGACCTCTGAATTCGTTGTCAAGATCTTG<br>CAATTCTTGGTTCCTATTCTAATTTTGGGTTTGGCTTTCGCTGTCAGACACTACACC<br>AAAGTTGAATAA |
| ClCPR | *Catharanthus longifolius* | [3] | **ATG**GATTCTTCTTCTGAAAAATTGTCTCCATTTGAATTGATGTCTGCTATTTTGAAAG<br>GTGCTAAATTGGATGGTTCTAATTCTTCTGATTCTGGTGTTGCTGTTTCTCCAGCTG<br>TTATGGCTATGTTGATGGAAAATAAAGAATTGGTTATGATTTTGACTACTTCTGTTG<br>CTGTTTTGATTGGTTGTGTTGTTGTTTTGATTTGGAGAAGATCTTCTGGTTCTGGTA<br>AAAAAGTTGTTGAACCACCAAAATTGATTGTTCCAAAATCTGTTGTTGAACCAGAAG<br>AAATTGATGAAGGTAAAAAAAAATTTACTATTTTTTTTGGTACTCAAACTGGTACTGC<br>TGAAGGTTTTGCTAAAGCTTTGGCTGAAGAAGCTAAAGCTAGATATGAAAAAGCTG<br>TTATTAAAGTTATTGATATTGATGATTATGCTGCTGATGATGAAGAATATGAAGAAAA<br>ATTTAAAAAAGAAACTTTGGCTTTTTTTATTTTGGCTACTTATGGTGATGGTGAACCA<br>ACTGATAATGCTGCTAGATTTTATAAATGGTTTGTCGAGGGAAACGACAGGGGAGA<br>CTGGCTGAAGAACTTACAGTACGGCGTCTTCGGATTAGGCAACCGTCAGTACGAG<br>CACTTCAACAAGATAGCAAAGGTCGTCGACGAGAAGGTGGCAGAGCAGGGCGGG<br>AAGAGGATCGTCCCCCTAGGACTAGGAGACGACGACCAGTGCATAGAGGACGACT<br>TCGCGGCGTGGCGTGAGAACGTCTGGCCCGAGCTAGACAACCTACTGAGGGACG<br>AGGACGACACAACCGTCTCAACCCCCTACACCGCAGCCATACCTGAGTACCGTTT<br>CGTATTCCACGACAAGTCGGATTCCTTAATAAGCGAGGCAAACGGACACGCAAAC<br>GGGTACGCAAACGGGAACACGGTATACGACGCCCAGCACCCCTGCAGGAGTAAC<br>GTCGCAGTCAGGAAGGAGCTACACACACCTGCCTCAGACCGTAGTTGCACGCACT<br>TAGAGTTCGACATCGCCGGCACAGGACTGTCCTACGGGACAGGAGACCACGTGG<br>GTGTGTACTGCGACAACTTATCCGAGACGGTAGAGGAAGCGGAGAGGCTACTTAA<br>CCTACCGCCTGAGACCTACTTCTCGTTACACGCAGACAAAGAGGACGGAACACCC<br>TTAGCAGGCAGCTCATTACCTCCTCCCTTCCCGCCTTGCACACTGAGGACAGCCC<br>TGACCAGGTACGCAGACTTATTAAACACGCCGAAGAAGAGCGCCCTACTAGCCTT<br>AGCCGCATACGCGTCAGACCCTAACGAGGCAAACAGGCTAAAGTACCTAGCCAGT |

| | | | |
|---|---|---|---|
| | | | CCGGCAGGAAAGGACGAGTACGCCCAGTCCCTGGTCGCCAACCAGCGTTCCTTAT<br>TAGAGGTAATGGCAGAGTTCCCCAGCGCAAAGCCCCCTCTGGGAGTATTCTTCGC<br>AGCAATAGCACCTCGTCTACAGCCTAGGTTCTACAGTATCTCATCAAGTCCGAGGA<br>TGGCAACATCAAGGATACACGTAACATGCGCGCTGGTATACGAGAAGACCCCGGG<br>CGGGAGAATACACAAGGGTGTATGCTCAACATGGATGAAGAACGCCATCCCCCTA<br>GAGGAGAGTCGTGACTGCTCCTGGGCACCTATATTCGTAAGGCAGAGCAACTTCA<br>AGTTACCTGCAGACCCTAAGGTACCTGTGATAATGATAGGTCCAGGTACAGGTTTG<br>GCTCCGTTTAGAGGTTTTCTTCAAGAAAGATTGGCCTTGAAAGAGGAAGGAGCAGA<br>ATTAGGTACTGCTGTCTTCTTCTTTGGTTGCAGGAACCGTAGGATGGACTACATAT<br>ACGAGGACGAGCTGAACCACTTCTTAGAGACAGGGGCATTATCCGAGTTATTAGTC<br>GCCTTCTCGCGTGAGGGCCCGACAAAGCAGTACGTGCAGCACAAGATGGCGGAG<br>AAGGCAAGCGACATATGGAGAATGATATCAGACGGAGCATACGTGTACGTCTGCG<br>GCGACGCAAAGGGAATGGCCAGGGACGTACACCGTACACTACACACCATCGCACA<br>GGAGCAGGGATCAATGGACTCCACCCAGGCTGAAGGTTTTGTTAAAAATTTGCAAA<br>TGACTGGTAGATATTTGAGAGATGTTTGGTAA |
| RsCPR | *Rauvolfia serpentina* | [3] | **ATG**GATTCTTCTTCTGAAAAATTGTCTCCATTTGAATTGATGTCTGCTATTTTGAAAG<br>GTGCTAAATTGGATGGTTCTAATTCTTCTGAATCTGGTGGTGCTGTTTCTCCAGCTG<br>TTGTTGCTATGTTGATGGAAAATAAAGAATTGGTTATGATTTTGACTACTTCTATGG<br>CTGTTTTGATTGGTTGTGTTGTTGTTTTGATGTGGAGAAGATCTTCTGGTTCTGCTA<br>AAAAAGTTGTTGAACCACCAAAATCTTTGGTTCCAAAGCTGTTGTTGAACCAGAAG<br>ATGTTGATGAAGGTAAAAAAAAATTTACTATTTTTTTTGGTACTCAAACTGGTACTGC<br>TGAAGGTTTTGCTAAAGCTTTGGCTGAAGAAGCTAAAGCTAGATATGAAAAAGCTA<br>TTATTAAAGTTATTGATATTGATGATTTTGCTGCTGATGATGAAGAATATGAAGAAAA<br>ATTGAAAAAAGAAACTTTGGCTTTTTTTATTTTGGCTACTTATGGTGATGGTGAACC<br>AACTGATAATGCTGCTAGATTTTATAAATGGTTTGTCGAGGGAAACGAGCGTGGAG<br>TATGGCTAAAGAACCTGCAGTACGGGGTATTCGGCCTTGGAAACAGGCAGTACGA<br>GCACTTCAACAAGATCGCGAAGGTAGTAGACGAGCAGTTAGCCGAGCAGGGCGG<br>AAAGCGTGTCGTACCGCTTGGCCTAGGCGACGACGACCAGTGCATAGAGGACGA<br>CTTCGCAGCCTGGAGAGAGACGGTATGGCCTGAGTTAGACCAGCTTTTAAGGGAC<br>GAGGACGACACCGCCGTAGCGACCCCGTACACCGCAGCAATCCCTGAGTACCGT<br>GTGGTCTTCCACCACAAGTCAGACTCGCTGATCTCCGAGGCAAACGGGCACGCGA<br>ACGGATACGCCAACGGACACACAGTATACGACGCACAGCACCCCTGCAGGTCAAA<br>CGTCGCAGTACGTAAGGAACTACACACACCCGCCTCGGAGAGGAGTTGCACACAC<br>TTAGAGTTCGACATCGCGGGAACGGGATTATCGTACGAGACGGGAGACCACGTAG<br>GGGTATACTGCGAGAACTTAACAGAGACGGTGGAAGAGGCGGAGAGGCTACTAAA<br>CTTACCCCCTGAGACCTACTTCTCCCTACACGCCGACAAGGAAGACGGGACGCCC<br>TTAGGTGGAAGCTCATTACCGCCACCTTTCCCGCCTTGCACCCTGCGTACCTCATT<br>AACCCAGTACGCAGACCTTCTGTCCACCCCGAAGAAGAGCGCATTACTGGCATTA<br>GCAAGCTACGCCAGCGACCCTAACGAGGCCGACCGTTTAAAGTACCTTGCCTCAC<br>CGGCCGGCAAGGACGAGTACGCGCAGAGCCTAGTCGCAAACCAGCGTAGTTTATT<br>AGAGGTCCTTGCGGAGTTCCCTTCCGCAAAGCCACCTCTTGGAGTATTCTTCGCA<br>GCCATCGCACCTAGGCTACAGCCTAGGTTCTACAGCATCAGCTCAAGTCCCAGGA<br>TGGCACCTAGCAGGATACACGTAACCTGCGCGCTGGTGTACGAGAAGACCCCTGG<br>CGGTAGGATACACAAGGGAGTGTGCAGTACCTGGATGAAGAACGCGATCCCTTTA<br>GAGGAGTCCCACGACTGCTCATGGGCACCCATCTTCGTAAGGCAGAGTAACTTCA<br>AGTTACCTACCGACCCTAAGGTCCCCATCTTAATGATAGGACCTGGCACGGGCCTT<br>GCACCGTTCAGGGGATTCTTACAGGAGAGGCTGGCATTAAAGGAAGAGGGCGCC<br>GAGCTGGGACCTGCCGTATTCTTCTTCGGCTGCAGGAACCGTAAGATGGACTTCA<br>TCTACGAGGACGAGCTAAACCACTTCCTTGAGACAGGAGCGGTGAGTGAGTTAAT<br>AGTGGCCTTCAGTAGGGAAGGACCTACAAAGCAGTACGTGCAGCACAAGATGGCC<br>GAGAAGGCTTCCGACATATGGAGAATGATAAGCGAGGGAGCCTACGTATACGTAT<br>GCGGCGACGCAAAGGGCATGGCCAGGGACGTCCACCGTACACTACACACAATCG<br>CACAGGAGCAGGGGTCAATGGACTCCAGCAAGGCTGAATCTTTGGTTAAATCTTTG<br>CAAATTTCTGGTAGATATTTGAGAGATGTTTGGTAA |
| AhCPR | *Amsonia hubrichtii* | | **ATG**GACTCTTCCAGTGAGAAATTGTCTCCATTTGATTTAATGACGGCTATATTAAGA<br>GGTGCCAAATTTGACGGGAGTAATAGTTCCGAGTCCGGGGGATTAGTCTCACCTG<br>CAGTAGTGGCAATGTTAATGGAGAACAAAGAGCTAATGATGATTCTAACCACGTCA<br>GTCGCCGTACTGATTGGGTGCGTCGTAGTCTTGATTTGGAGGCGTTCAAGTGGTA<br>GTGCGAAAAAAGTAGTTGAACCTCCAAAGCTGCAAGTCCCCAAAGTGGACGTGGA |

| | | | |
|---|---|---|---|
| | | | GCCCGAGGAGGTCGACGACGGCAGTAAAAAAGTGACGATCTTCTTTGGGACTCAA<br>ACAGGTACAGCCGAGGGCTTCGCCAAAGCGCTAGCCGAGGAGGGTAAAGCTAGG<br>TATGAAAAAGCAACGTTTAAGGTCATTGACTTGGACGATTACGCAGCGGACGACGA<br>AGAGTACGAAGAGAAATTAAAGAAAGAGACCTTAGCGTTTTTTTTCCTGGCCACGT<br>ACGGTGATGGGGAGCCTACGGACAACGCCGCGAGATTCTATAAATGGTTTGCGGA<br>AGGGAAGGAGAGAGGTGATTGGCTGAAGAACTTACAATACGGGGTGTTCGGCTTA<br>GGTAATCGTCAGTACGAGCATTTTAATAAGATTGCCAAAGTTGTGGACGAGTTGGT<br>GGCTGAGCAGGGGGGAAAGCGTTTAGTCCCACTAGGTCTAGGCGACGATGACCA<br>ATGTATTGAAGACGACTTCGCTGCGTGGAGGGAAAATGTGTGGCCCGAGCTAGAT<br>AAACTATTACGTGATGAAGACGATACTACAGTGTCTACACCATATACAGCAGCTATC<br>TTGGAGTACAGGGTAGTGATTCATGATAGATCTGATACCTTGATAAGTGAAGCTAA<br>CGGGCACGCTAACGGGTACGCTAATGGTAATACTGTCTACGACGCGCAACATCCC<br>TGCAGGAGCAACGTTGCCGTTAAGAAAGAATTGCACACACCAGCAAGTGATCGTT<br>CTTGTACTCACCTTGAGTTTGATATCGCCGGAACCGGGCTTTCTTACGAAACAGGC<br>GATCACGTCGGCGTGTACTGCGAAAATCAGATCGAAACTGTAGAAGAAGCTGAGC<br>GTATTTTAAACCTTCCGCCTGACACATATTTTAGCATACACACGGATAAGGAGGAC<br>GGCACTAAACTTGGCGGGAGTTCTCTACAACCACCCTTTCCGCCTTGTACATTAAG<br>GACTGCCTTAACAAGATACGCCGATTTGCTGTCAAGCCCGAAAAAGAGTGCCCTTC<br>TAGCCCTAGCTGCCTACGCATCAGACCCAAAGGAAGCCGATCGTTTAAAGTATTTA<br>GCCAGCCCGGCTGGCAAAGATGAATATGCACAATGGTTAGTAGCCAATCAGAGAT<br>CTTTGCTAGAAGTTATGGCAGAATTTCCTAGTGCTAAGCCCCCCTTAGGGGTGTTC<br>TTTGCGGCCGTCGCCCCACGTCTTCAGCCCAGGTTCTATTCTATTAGTTCTAGTCC<br>CCGTATGGCCCCATCCCGTATACATGTGACCTGCGCGCTTGTGTATGAAAAAACAC<br>CCACGGGGAGAATACACAAGGGAGTCTGTAGCACATGGATGAAGAATGCGATACC<br>CCTTGAAGAGTCTCGTGATTGTAACTGGGCGCCAATCTTCGTTAGACAAAGCAACT<br>TCAAGCTGCCCGCGGATCCAAAAGTCCCCATTATTATGATCGGGCCCGGAACAGG<br>GCTGGCTCCATTTAGAGGGTTCTTACAAGAAAGACTTGCTCTTAAGGAAGAAGGGG<br>CTGAACTAGGGCCCGCCGTATTTTACTTTGGTTGTAGAAACCGTAAGTTGGATTAC<br>ATTTACGAGGATGAGCTTAATCATTTCTTGGAGACCGGGGCCGTGTCTGAATTAGT<br>GGTAGCATTCAGTAGAGAAGGTCCAACTAAACAGTATGTACAACATAAGATGGCAG<br>AGAAGGCTTCCAATATATGGAGACTTATATCCGAAGGCGCCTATGTTTACGTTTGC<br>GGGGACGCTAAGGGCATGGCGAGAGACGTGCACAGGACGCTTCACACAATCGCG<br>CAAGAACAAGGATCTATGGACAGCACGAAGGCTGAGGGATTTGTTAAAAATCTTCA<br>GATGAACGGCAGGTACCTTAGAGATGTGTGGTAA |
| AniCPR | *Aspergillus niger* | [3] | **ATG**GCTCAATTGGACACCCTTGACCTAGTCGTGCTTGCGGTACTTCTGGTCGGCT<br>CCGTCGCATATTTTACCAAAGGGACATATTGGGCAGTTGCTAAAGATCCGTATGCT<br>TCAACAGGACCTGCTATGAACGGCGCGGCGAAGGCAGGTAAGACTCGTAACATCA<br>TCGAGAAGATGGAGGAAACCGGTAAAAATTGCGTAATATTCTACGGGTCTCAAACT<br>GGGACTGCTGAGGACTACGCGTCCCGTCTGGCAAAAGAAGGAAGTCAGAGATTTG<br>GCCTAAAAACCATGGTGGCAGACTTGGAAGAGTACGATTACGAGAACCTTGACCA<br>GTTTCCCGAAGACAAGGTGGCTTTCTTTGTCTTAGCTACTTATGGAGAAGGGGAAC<br>CTACTGATAATGCTGTCGAGTTCTATCAGTTTTTTACGGGCGATGATGTGGCGTTT<br>GAGTCCGCGAGCGCTGATGAAAAGCCTCTGTCAAAGCTAAAATACGTCGCATTTG<br>GTCTAGGGAATAACACCTATGAACACTACAACGCGATGGTCAGGCAGGTTGACGC<br>AGCATTTCAGAAGCTAGGACCGCAGCGTATCGGTTCAGCAGGGGAGGGGGATGA<br>CGGAGCCGGAACAATGGAGGAGGATTTTCTGGCGTGGAAAGAGCCAATGTGGGC<br>TGCTCTTTCCGAGTCTATGGATCTAGAGGAAAGGGAAGCTGTATATGAACCGGTTT<br>TTTGCGTCACAGAAAATGAGAGCCTTAGTCCTGAAGATGAAACCGTATACCTGGGT<br>GAACCTACACAATCACACTTACAGGGTACGCCTAAAGGTCCGTATAGCGCTCACAA<br>CCCCTTTATTGCTCCAATCGCGGAATCAAGAGAACTATTCACTGTTAAGGACCGTA<br>ACTGTCTACACATGGAGATTTCAATAGCAGGATCTAATCTAAGTTACCAAACGGGA<br>GATCATATTGCAGTCTGGCCGACCAATGCCGGTGCGGAAGTCGATAGGTTCTTAC<br>AGGTTTTTGGTCTTGAAGGCAAAAGAGACAGCGTGATCAACATTAAAGGAATAGAC<br>GTAACGGCGAAAGTTCCGATCCCTACACCAACGACGTATGACGCAGCAGTGAGAT<br>ATTATATGGAGGTCTGTGCGCCGGTCTCCAGACAGTTCGTGGCGACGTTGGCAGC<br>TTTCGCTCCAGACGAAGAATCTAAAGCGGAGATCGTCAGGTTAGGATCACACAAG<br>GACTACTTCCACGAAAAAGTCACTAACCAATGCTTTAATATGGCTCAAGCACTGCA<br>GTCCATCACCTCAAAGCCGTTTAGTGCTGTACCTTTCAGTCTGTTGATTGAGGGAA<br>TTACGAAGCTTCAGCCTAGATATTACTCAATCTCCTCATCTTCACTAGTCCAGAAAG |

| | | | |
|---|---|---|---|
| | | | ATAAGATATCTATCACTGCCGTCGTCGAATCTGTTAGGTTACCAGGTGCTTCTCAC<br>ATGGTTAAAGGAGTGACGACCAACTATTTGTTAGCACTAAAACAAAAACAAAACGG<br>CGATCCCTCACCGGACCCTCACGGGCTAACTTATAGCATTACAGGCCCGAGAAAT<br>AAGTACGATGGGATACATGTCCCAGTGCACGTGAGGCACAGTAACTTTAAGTTACC<br>CTCTGACCCATCTAGACCTATAATTATGGTTGGGCCGGGAACGGGTGTGGCGCCC<br>TTTAGAGGGTTCATACAAGAAAGAGCGGCTCTAGCAGCGAAGGGAGAAAAGGTAG<br>GACCTACTGTACTGTTTTTTGGTTGCCGTAAAAGTGACGAGGACTTCCTATATAAG<br>GATGAATGGAAAACGTATCAAGACCAGTTAGGAGACAATCTGAAGATTATAACCGC<br>CTTTAGTAGGGAAGGACCTCAGAAAGTCTACGTACAACACCGTCTAAGGGAACACT<br>CCGAACTAGTATCCGATCTTCTTAAACAAAAGGCAACCTTTTATGTTTGTGGTGACG<br>CGGCTAACATGGCGAGAGAAGTTAATTTAGTACTAGGACAAATAATTGCAGCTCAA<br>AGAGGACTACCGGCAGAGAAGGGTGAGGAGATGGTGAAGCACATGAGGCGTCGT<br>GGGAGGTATCAAGAAGACGTCTGGAGTTAA |
| OeCPR | *Olea europaea* | [3] | **ATG**GACTCCACCTCAGAGAAGCTTTCACCGTTCGACTTCCTGACGGCAATCCTGAA<br>GGGAGTGAAGGTGGACCAGAGTAACGGCAGTTTAGACGTACCCCCTGCATTAGCC<br>AAGATATTAATGGAGAACAGGGACCTGATGATGGTATTAACAACATCAGTGGCCTT<br>ACTAATCGGATGCGTAGTAGTCCTTGTATGGAGACGTGCAGCAGGAAGTGCAAAG<br>AAGTTAGTCGAGCCGCCTAAGTTAGTAATACCTAAGGCCGTAGCCGAGTTAGAAGA<br>GGTGGACGACGGAAAGAAGAAGGTCACCATATTCTTCGGGACCCAGACAGGGAC<br>CGCCGAGGGATTCGCAAAGGCCTTAGCCGAGGAAGGGAACGCAAGGTACGAGAA<br>GGCAACCTTCAAGGTCATAGACTTAGACGACTACGCAGCCGACGACGAGGAGTAC<br>GAGGAGAAGTTAAAGAAGGAGACATTAGCCTTCTTCTTCCTTGCAACATACGGAGA<br>CGGAGAGCCTACAGACAACGCCGCCAGGTTCTACAAGTGGTTTGCCGAGGGAAA<br>GGAGCGTGGAGACTGGTTAAAGAACCTGCAGTACGGCGTCTTCGGATTAGGCAAC<br>AGGCAGTACGAGCACTTCAACAAGATCGCAAACGTCGTCGACGAGCTTATCGCAA<br>AGCAGGGCGGAAAGCAGCTGATACCGGTAGGCCTAGGGGACGACGACCAGTGCA<br>TAGAGGACGACTTCGCCGCCTGGCGTGAGCTAGTGTGGCCTGAGCTTGACCAGTT<br>ACTAAGGGACGAGGACGACGCGTCGGCAGCAACGCCGTACACGGCAGCAGTCCT<br>TGAGTACAGGGTCGTATTCCACGACAAGCCTGACGCGTCATTAAGCGAGAACGGA<br>TCCATACACGCAAACGGGCACGCGGTGTACGACGCGCAGCACCCTTGCAGGGCA<br>AACGTAGCCGTGAAGAGGGAGTTACACTCGCCGGCATCCGACAGGTCATGCACCC<br>ACCTAGAGTTCGACATATCCGGAACAGGACTATCCTACGAGACAGGAGACCACGT<br>CGGCGTGTACTGCGAGAACCTGATCGAGGTCGTGGAAGAGGCAGAGCAGTTACTA<br>AACATACCACCCGACACATACTTCTCCATCCACACAGAGAAAGAGGACGGCACGC<br>CCTTATCAGGCTCAAGCCTACCCCCACCTTTCCCGCCTTGCACATTACGTACCGCG<br>TTATCAAGGTACGCAGACTTATTAAGTGCGCCCAAGAAGAGCGCGCTAATCGCGC<br>TTGCGACATACGCGTCCGACCCTAGTGAGGCAGACCGTCTAAAGTACTTAGCGAG<br>TCCTTCAGGAAAAGAGGAGTACGCGCAGTGGATAGTCGCCAACCAGAGGTCCTTA<br>CTAGAGGTAATGGCAGAGTTCCCTTCCGCAAAGCCACCCCTTGGCGTCTTCTTCG<br>CAGCAGTCGCACCCCGTCTTCAGCCTAGGTTCTACAGTATCAGTTCCTCACCTAAG<br>ATCGCACCGAGTAGGATCCACGTGACCTGCGCGCTGGTATACGAGAAGACACCTG<br>CAGGCAGGATCCACAAAGGGGTATGCTCCACATGGATGAAGAACGCCCTGCCCCT<br>GGAGCAGTCACCTGACTGCTCCTGGGCGCCCATATTCGTACGTAACTCCAACTTC<br>AAGCTACCTGCCGACCCTAAGGTGCCTATCATAATGATCGGACCGGGAACAGGCC<br>TAGCCCCCTTCCGTGGCTTCTTACAGGAGAGGCTGGCACTAAAGGAGAGCGGAGC<br>CGAGCTGGGCCCTGCGGTCTTATTCTTCGGATGCCGTAACTCAAAGATGGACTAC<br>ATCTACGAGGACGAGTTAAAGAACTTCTTCCAGGACGGAATAATATCGGAGCTGGT<br>AGTCGCCTTCTCAAGGCAGGGACCTACAAAGGAGTACGTACAGCACAAGATGACA<br>GAGAAGGCCTCAAACATATGGAACATGTTAAGCGAGGGAGCCTACATCTACGTGT<br>GCGGAGACGCCAAGGGAATGGCAAGGGACGTACACAGGACCCTACACACAATAG<br>CACAGGAGCAGGGTGCATTAGACTCCTCAAAGGCTGAATCTATGGTTAAAAATTTG<br>CAAATGACTGGTAGATATTTGAGAGATGTTTGGTAA |
| CpCPR | *Chrysomela populi* | [3] | **ATG**GATGAAACTGAAGCTACTTTGGGTACTCAACAAATTTCTGAAGCTTCTGATTTT<br>TTGTTTTTGCCATTGGATATTTTGATTGTTGCTTTGATTGTTGGTGGTATTTCTTGGT<br>ATTTGTTGTCTAAACATAAAAAAACTACTACTGTTTCTTCTGGTAGATCTTATTCTAT<br>TCAACCAACTTCTATTGCTTTGCAAAATCCATCTGATAATTCTTTTATTAAAAAATTG<br>CAAACTTCTGGTAGATCTTTGGTTGTTTTTTATGGTTCTCAAACTGGTACTGGTGAA<br>GAATTTGCTGGTAGATTGGCTAAAGAAGGTGCTAGATATAGATTGAAAGGTATGGT<br>TGCTGATCCAGAAGAATGTGATATGGAAGAATTGGTTAATTTGAAAAATATTCCAAA |

| | | | |
|---|---|---|---|
| | | | TTCTTTGGCTGTTTTTTGTTTGGCTACTTATGGTGAAGGTGATCCAACTGATAATGC<br>TATGGAATTTTATGAATGGTTGCAAAATGGTGATGCTGATTTGTCTGGTTTGACTTA<br>TGCTGTTTTTGGTTTGGGTAATAAAACTTATGAACATTATAATGAAGTTGCTATTTAT<br>GTTGATAAAAGATTGGAAGAATTGGGTGCTACTAGAGTTTATGATTTGGGTTTGGG<br>TGATGATGATGCTAATATTGAAGATGATTTTATTACTTGGAAAGATAAATTTTGGCC<br>AGCTGTTTGTGAACATTTTGGTATTGAAGCTACTGGTGAAGATATTAATATGCATCA<br>ATATAGATTGGAAGAATTTGAAGAAGTTCCAGATAGAGTTTTTTTTGGTGAAATGGC<br>TAGATTGCATTCTTTGAAAAATCAAAGACCACCATATGATGCTAAAAATCCATATTT<br>GGCTACTATTAAAGTTAATAGAGAATTGCATCAACAAGGTGATAGATCTTGTATGCA<br>TATTGAATTGGATATTGAAGGTTCTAAAATGAGATATGAATCTGGTGATCATTTGGC<br>TATTTATCCAGTTAATGATGCTGCTTTGGTTGAAAAATTGGGTAGATTGTGTGGTAA<br>AGATTTGGAAACTATTTTTATGTTGATTAATACTGATGAAGAATCTTCTAAAAAACAT<br>CCATTTCCATGTCCATGTTCTTATAGAACTGCTTTGACTCATTATTTGGATATTACTC<br>AAAATCCAAGAACTCATGTTTTGAAAGAATTGGCTGAATATTGTTCTGATGAAGCTG<br>CTAAAGCTAAATTGATGTTGATGTCTTCTACTTCTCCAGAAGGTAAAGCTTTGTATC<br>AATCTTGGATTATTGAAGATAATAGAAATATTGTTCATGTTATTGAAGATACTCCATC<br>TTGTCAACCAGCTTTGGATCATTTGTGTGAATTGTTGCCAAGATTGCAACCAAGATA<br>TTATTCTATTTCTTCTTCTCCAAAATTGTATCCAAATACTGTTCATATTACTGCTGTT<br>GTTGTTGAATATGATACTCCAACTGGTAGACATAATAAAGGTGTTGCTACTACTTGG<br>TTGAAAACTAAAATTCCAGATCCAGATAAAGAACCAATGTTGGTTCCAGTTTTTATT<br>AGAAAATCTCAATTTAGATTGCCAATTAAACCACAAACTCCAATTATTATGATTGGTC<br>CAGGTACTGGTTTGGCTCCATTTAGAGGTTTTATTCAAGAAAGAGATTTGATTAGAT<br>CTGAAGATAAATCTGTTGGTGAAACTATTTTGTATTTTGGTTGTAGAAAAAGAAAAG<br>AAGATTTTTTGTATGGTGATGAATTGTTGGGTTATGAAAAATCTGGTTCTTTGACTTT<br>GCATTTGGCTTTTTCTAGAGATCAACAACAAAAAGTTTATGTTACTCATTTGTTGCAA<br>CAACATGCTGAAGAAGTTTGGAGAGTTATTGGTGAAAATTCTGGTCATGTTTATATT<br>TGTGGTGATGCTAAAACTATGGCTTTTGATGTTAGATCTATTTTGGCTAAAATTTTG<br>CAAGAAAAAGGTCAAATGACTGAACAACAAGCTTTGGCTTATTTGAAAAAAATGGAA<br>ACTCAAAAAAGATATTCTGCTGATGTTTGGTCTTAA |
| CrCPR | *Catharanthus roseus* | [3] | **ATG**GACTCATCCTCCGAGAAGTTGTCACCATTCGAACTTATGTCAGCAATTCTTAA<br>GGGAGCCAAGCTGGACGGTAGTAACAGTTCTGATTCCGGTGTCGCTGTATCACCT<br>GCTGTTATGGCAATGTTACTAGAAAATAAAGAGTTAGTAATGATATTGACGACATCT<br>GTCGCTGTCTTGATTGGTTGCGTCGTTGTGCTAATTTGGCGTAGATCTTCAGGGTC<br>CGGTAAGAAGGTTGTGGAGCCACCCAAGTTGATAGTCCCAAAAAGTGTAGTGGAG<br>CCAGAAGAAATAGATGAAGGAAAAAAAAAATTCACTATCTTCTTTGGTACACAAACT<br>GGGACAGCTGAAGGTTTTGCTAAGGCTTTAGCCGAAGAAGCAAAGGCTAGATACG<br>AAAAGGCAGTTATAAAAGTAATCGATATTGACGATTATGCAGCAGACGATGAGGAG<br>TATGAGGAAAAATTCAGAAAAGAGACTTTGGCCTTCTTTATATTGGCAACATATGGC<br>GATGGTGAGCCTACTGATAACGCTGCAAGGTTTTACAAATGGTTTGTAGAGGGTAA<br>TGATAGAGGTGACTGGCTTAAGAACTTACAGTATGGCGTCTTCGGTTTGGGCAATA<br>GACAGTATGAACATTTCAATAAGATTGCAAAAGTTGTAGATGAAAAGGTTGCCGAG<br>CAAGGAGGGAAGAGGATAGTGCCTTTAGTTTTAGGAGACGATGATCAATGTATTGA<br>AGATGACTTTGCTGCATGGAGAGAAAACGTCTGGCCTGAACTGGATAATCTGCTAA<br>GAGACGAGGATGATACTACAGTGTCTACTACCTATACAGCCGCTATACCAGAATAC<br>AGAGTTGTTTTCCCTGATAAAAGTGATTCTTTGATTTCTGAGGCCAACGGCCACGC<br>TAACGGCTATGCGAACGGCAATACTGTATACGATGCTCAACACCCTTGCCGTAGTA<br>ACGTCGCTGTCAGGAAAGAGTTACATACCCCAGCTTCTGATAGGTCTTGTACACAT<br>TTGGATTTTGATATAGCGGGTACTGGATTATCATATGGTACAGGAGATCACGTCGG<br>TGTCTATTGTGACAATTTATCTGAGACCGTAGAAGAAGCAGAAAGGTTGTTGAACTT<br>GCCCCCTGAAACGTATTTCAGTTTGCACGCTGATAAAGAAGATGGTACTCCATTAG<br>CAGGATCATCATTGCCCCCTCCTTTTCCTCCGTGCACATTGAGGACTGCCCTTACT<br>AGATATGCTGATCTGCTGAATACCCCAAAAAAGTCCGCCTTGCTGGCTCTAGCTGC<br>TTATGCAAGCGATCCAAATGAGGCTGATCGTTTGAAGTACTTGGCAAGCCCAGCTG<br>GCAAAGACGAGTATGCTCAATCTTTGGTAGCTAATCAGAGAAGCCTGCTAGAAGTA<br>ATGGCTGAATTTCCTTCCGCCAAGCCACCGTTAGGTGTATTCTTCGCAGCAATAGC<br>TCCCAGATTACAACCCAGATTCTACTCTATATCTTCTAGCCCAAGGATGGCCCCTT<br>CCAGAATTCATGTCACTTGCGCTCTAGTTTATGAGAAAACTCCAGGCGGGAGGATT<br>CATAAAGGCGTATGTTCAACTTGGATGAAGAACGCTATTCCTCTGGAGGAATCTCG<br>TGATTGTAGCTGGGCACCGATCTTTGTGAGACAGTCTAACTTTAAGCTGCCTGCCG |

| | | | |
|---|---|---|---|
| | | | ATCCAAAAGTTCCTGTAATCATGATAGGTCCAGGCACCGGGCTAGCACCTTTTAGA<br>GGTTTCCTTCAAGAAAGACTTGCTCTGAAAGAGGAAGGAGCTGAATTAGGAACTGC<br>TGTATTTTTTTTTGGGTGTAGGAACAGAAAAATGGATTATATATATGAAGATGAATT<br>GAATCATTTCTTGGAAATCGGCGCGTTATCAGAATTGCTGGTTGCATTCAGTAGGG<br>AAGGTCCTACTAAGCAATATGTTCAACACAAAATGGCCGAAAAAGCCAGTGATATT<br>TGGCGTATGATCTCTGATGGTGCTTATGTTTATGTCTGCGGAGATGCCAAGGGCAT<br>GGCCAGAGACGTTCATAGGACATTACACACTATAGCTCAAGAGCAAGGATCAATG<br>GACTCTACTCAGGCCGAAGGATTTGTGAAAAACTTACAAATGACCGGTAGATATTT<br>AAGAGACGTATGGTAA |
| AnnCPR | *Artemisia annua* | [3] | **ATG**CAATCTACAACCTCCGTCAAGTTATCACCCTTTGACCTGATGACCGCATTACTA<br>AACGGTAAAGTCTCTTTTGATACTAGTAATACTTCCGATACAAACATTCCATTAGCT<br>GTCTTCATGGAAAATAGAGAGCTGTTGATGATTTTGACCACATCAGTCGCTGTTTTG<br>ATTGGGTGCGTAGTAGTTTTGGTGTGGAGAAGATCTTCTTCCGCGGCGAAAAAAG<br>CAGCTGAATCTCCCGTTATAGTTGTTCCAAAAAAAGTTACCGAAGACGAAGTGGAT<br>GACGGTAGAAAAAAGGTCACTGTTTTTTTTGGTACACAGACAGGCACTGCAGAAGG<br>CTTCGCTAAGGCACTAGTTGAAGAGGCCAAAGCAAGGTATGAAAAAGCCGTATTCA<br>AGGTTATAGATCTGGATGATTATGCTGCCGAGGATGATGAATACGAAGAGAAGTTA<br>AAAAAGGAGTCATTGGCATTTTTTTTCTTAGCAACCTACGGAGACGGCGAGCCTAC<br>CGACAATGCTGCCCGTTTTTATAAATGGTTCACTGAAGGTGAAGAAAAGGGAGAAT<br>GGTTAGATAAATTACAATATGCAGTCTTCGGCCTTGGGAATAGACAGTACGAGCAT<br>TTTAACAAGATCGCCAAAGTTGTAGATGAAAAACTGGTCGAACAAGGAGCTAAGCG<br>TTTAGTGCCAGTTGGCATGGGTGATGATGACCAGTGTATCGAAGACGACTTTACGG<br>CTTGGAAAGAATTGGTATGGCCAGAACTAGATCAATTGTTGAGAGATGAGGATGAT<br>ACATCCGTCGCAACACCTTACACTGCTGCAGTGGGAGAATACAGAGTAGTTTTCCA<br>CGATAAACCAGAAACCTACGATCAGGACCAATTAACAAATGGACATGCTGTTCATG<br>ATGCTCAACACCCTTGCAGATCTAATGTCGCTGTTAAGAAAGAACTTCACTCTCCA<br>CTAAGCGATAGATCATGCACTCATCTAGAATTTGATATCTCAAATACAGGTCTTTCT<br>TATGAAACTGGTGATCATGTTGGTGTATATGTTGAAAATCTTAGTGAGGTAGTGGA<br>CGAAGCTGAAAAATTAATAGGCTTGCCTCCACATACGTACTTCTCTGTTCATACTGA<br>TAACGAGGATGGTACACCACTTGGTGGTGCTAGTTTACCACCTCCATTTCCGCCAT<br>GTACATTAAGGAAAGCGTTGGCTAGTTATGCGGATGTTCTTTCTAGTCCTAAAAAAT<br>CCGCTTTATTGGCTCTAGCTGCACATGCAACCGACAGTACTGAAGCTGATAGACTG<br>AAATTCTTCGCGTCTCCAGCAGGAAAGGACGAATACGCCCAATGGATTGTCGCTA<br>GTCATAGATCATTACTAGAAGTGATGGAAGCATTCCCGTCTGCCAAGCCCCCTTTG<br>GGCGTATTTTTTGCTTCCGTAGCGCCTAGATTACAACCAAGATACTACTCCATTTCT<br>TCATCACCTAAATTCGCGCCAAATAGAATTCATGTAACGTGCGCTTTGGTCTACGA<br>ACAAACACCTTCCGGTCGTGTTCACAAAGGAGTATGTAGTACCTGGATGAAAAACG<br>CAGTTCCTATGACGGAATCCCAGGACTGTTCATGGGCTCCAATTTATGTCAGAACG<br>AGTAACTTTAGGTTACCCAGCGACCCAAAGGTTCCAGTCATAATGATAGGCCCAGG<br>TACAGGATTGGCTCCCTTTAGAGGATTTCTACAAGAGAGACTTGCGCAAAAAGAAG<br>CAGGTACGGAACTGGGAACTGCCATTCTATTTTTTTGGTTGCAGGAATAGAAAGGTG<br>GATTTCATTTACGAAGATGAACTTAATAATTTCGTGGAAACTGGCGCCCTGTCAGA<br>GTTGGTCACGGCTTTCTCAAGAGAAGGAGCTACCAAGGAATACGTTCAACATAAGA<br>TGACTCAGAAAGCTTCTGATATCTGGAACCTGCTGTCCGAAGGAGCTTACTTGTAT<br>GTGTGTGGGGACGCAAAAGGTATGGCGAAGGATGTGCATAGAACCCTTCACACCA<br>TAGTTCAAGAACAAGGCAGTTTAGATTCTAGTAAAGCAGAATTGTACGTTAAGAACC<br>TGCAGATGGCTGGTAGATACTTAAGAGATGTTTGGTAA |

#### Supplementary table 2. Plasmids cloned and used in this study

| Name | Description | Reference |
|---|---|---|
| pCfB9336 | pgRNA_II-1_NatMX | [5] |
| pCfB9337 | pgRNA_IV-1_NatMX | [5] |
| pCfB9355 | II-1_Markerfree_BackBone | [5] |
| pCfB9356 | IV-1_Markerfree_BackBone | [5] |
| MoClo Yeast Toolkit (YTK) | https://www.addgene.org/kits/moclo-ytk/ | [6] |
| pMHGG2_Vec_II-1 | preassembled yeast MoClo level-2 backbone with GFP dropout for genomic integration in site II-1 | [7] |
| pMHGG0_3_NtCYP96T6 | Level-0 yeast MoClo plasmid containing NtCYP96T6 CDS part. | This study. |
| pMHGG0_3_NtNMT1 | Level-0 yeast MoClo plasmid containing NtNMT1 CDS part. | This study. |
| pMHGG0_3_NtAKR1 | Level-0 yeast MoClo plasmid containing NtAKR1 CDS part. | This study. |
| pMHGG1_Vec_Pos1 | Level-1 yeast MoClo vector plasmid containing GFP dropout. Includes 2µ yeast ORI and URA yeast selection marker. Gene position number 1 in level-2 plasmid. | [7] |
| pMHGG1_Vec_Pos2 | Level-1 yeast MoClo vector plasmid containing GFP dropout. Includes 2µ yeast ORI and URA yeast selection marker. Gene position number 2 in level-2 plasmid. | [7] |
| pMHGG1_Vec_Pos3* | Level-1 yeast MoClo vector plasmid containing GFP dropout. Includes 2µ yeast ORI and URA yeast selection marker. Gene position number 3 and final gene in level-2 plasmid. | [7] |
| pMHGG1_NtCYP96T6_Pos1 | Level-1 yeast MoClo plasmid containing pTDH3-NtCYP96T6-tTDH1 transcription unit. Includes 2µ yeast ORI and URA yeast selection marker. | This study. |
| pMHGG1_NtNMT1_Pos2 | Level-1 yeast MoClo plasmid containing pTEF1-NtNMT1-tENO2 transcription unit. Includes 2µ yeast ORI and URA yeast selection marker. | This study. |
| pMHGG1_NtAKR1_Pos3* | Level-1 yeast MoClo plasmid containing pCCW12-NtAKR1-tPKG1 transcription unit. Includes 2µ yeast ORI and URA yeast selection marker. | This study. |
| pMHGG2_NtCYP96T6-NtAKR1-NtNMT1@II-1 | Level-2 multigene vector for integration at site II-1. Contains transcription units for NtCYP96T6, NtNMT1 and NtAKR1. | This study. |
| pBJL87 | IV-1 integration plasmid for pTEF1-ClCPR-tCYC1 and pPGK1-CrCYB5-tVPS13. | [3] |
| pBJL88 | IV-1 integration plasmid for pTEF1-RsCPR-tCYC1 and pPGK1-CrCYB5-tVPS13. | [3] |
| pBJL89 | IV-1 integration plasmid for pTEF1-AhCPR-tCYC1 and pPGK1-CrCYB5-tVPS13. | [3] |
| pBJL90 | IV-1 integration plasmid for pTEF1-AniCPR-tCYC1 and pPGK1-CrCYB5-tVPS13. | [3] |
| pBJL91 | IV-1 integration plasmid for pTEF1-OeCPR-tCYC1 and pPGK1-CrCYB5-tVPS13. | [3] |

| | | |
|---|---|---|
| pBJL92 | IV-1 integration plasmid for pTEF1-CpCPR-tCYC1 and pPGK1-CrCYB5-tVPS13. | [3] |
| pBJL93 | IV-1 integration plasmid for pTEF1-CrCPR-tCYC1 and pPGK1-CrCYB5-tVPS13. | [3] |
| pBJL94 | IV-1 integration plasmid for pTEF1-AnnCPR-tCYC1 and pPGK1-CrCYB5-tVPS13. | [3] |
| pBJL95 | IV-1 integration plasmid for pTEF1-AtATR1-tCYC1 and pPGK1-CrCYB5-tVPS13. | [3] |
| pMHO46 | IV-1 integration plasmid for pTEF1-AtATR1-tCYC1. | This study. |
| pMHO47 | IV-1 integration plasmid for pTEF1-AtATR1-tCYC1 and pPGK1-NpCYB5-tVPS13. | This study. |
| pMHO48 | IV-1 integration plasmid for pTEF1-NpCPR-tCYC1 and pPGK1-CrCYB5-tVPS13. | This study. |
| pMHO49 | IV-1 integration plasmid for pTEF1-NpCPR-tCYC1. | This study. |
| pMHO50 | IV-1 integration plasmid for pTEF1-NpCPR-tCYC1 and pPGK1-NpCYB5-tVPS13. | This study. |
| pMHO51 | IV-1 integration plasmid for pTEF1-LrCPR2-tCYC1 and pPGK1-CrCYB5-tVPS13. | This study. |
| pMHO52 | IV-1 integration plasmid for pTEF1-LrCPR2-tCYC1. | This study. |
| pMHO53 | IV-1 integration plasmid for pTEF1-LrCPR2-tCYC1 and pPGK1-NpCYB5-tVPS13. | This study. |
| pMHO42 | CEN-ARS plasmid with URA marker for expression of 4NB2 RamR biosensor variant for 4OMe-norbelladine in yeast. | This study. |
| pCCD3 | Yeast genome integration plasmid of RamR reporter construct pCCW12-1X_RamO-yeGFP-tCPS1 at site XI-3. | [8] |

#### Supplementary table 3. Strains used in this study

| Name | Parent | Genotype | Reference |
|---|---|---|---|
| MIA-B0 | CEN.PK2-1C | MATa; his3D1; leu2-3_112; ura3-52; trp1-289; pTEF1-SpCas9-tCYC1. | [9] |
| yCCD7 | MIA-B0 | MATa; his3D1; leu2-3_112; ura3-52; trp1-289; PTEF1-SpCas9-TCYC1, **PCCW12_1XRamO-yeGFP-TCPS1**. | [8] |
| yMHO115 | yCCD7 | MATa; his3D1; leu2-3_112; ura3-52; trp1-289; PTEF1-SpCas9-TCYC1, PCCW12_1XRamO-yeGFP-TCPS1, **pMHO42.** | This study. |
| yCCD45 | MIA-B0 | MATa; his3D1; leu2-3_112; ura3-52; trp1-289; pTEF1-SpCas9-tCYC1, **pTDH3-NtCYP96T6-tTDH1, pTEF1-NtNMT1-tENO2, pCCW12-NtAKR1-tPKG1**. | This study. |
| yMHO52 | yCCD45 | MATa; his3D1; leu2-3_112; ura3-52; trp1-289; pTEF1-SpCas9-tCYC1, pTDH3-NtCYP96T6-tTDH1, pTEF1-NtNMT1-tENO2, pCCW12-NtAKR1-tPKG1, pTEF1-ClCPR-tCYC1, pPGK1-CrCYB5-tVPS13. | This study. |
| yMHO53 | yCCD45 | MATa; his3D1; leu2-3_112; ura3-52; trp1-289; pTEF1-SpCas9-tCYC1, pTDH3-NtCYP96T6-tTDH1, pTEF1-NtNMT1-tENO2, pCCW12-NtAKR1-tPKG1, pTEF1-RsCPR-tCYC1, pPGK1-CrCYB5-tVPS13. | This study. |
| yMHO54 | yCCD45 | MATa; his3D1; leu2-3_112; ura3-52; trp1-289; pTEF1-SpCas9-tCYC1, pTDH3-NtCYP96T6-tTDH1, pTEF1-NtNMT1-tENO2, pCCW12-NtAKR1-tPKG1, pTEF1-AhCPR-tCYC1, pPGK1-CrCYB5-tVPS13. | This study. |
| yMHO55 | yCCD45 | MATa; his3D1; leu2-3_112; ura3-52; trp1-289; pTEF1-SpCas9-tCYC1, pTDH3-NtCYP96T6-tTDH1, pTEF1-NtNMT1-tENO2, pCCW12-NtAKR1-tPKG1, pTEF1-AniCPR-tCYC1, pPGK1-CrCYB5-tVPS13. | This study. |
| yMHO56 | yCCD45 | MATa; his3D1; leu2-3_112; ura3-52; trp1-289; pTEF1-SpCas9-tCYC1, pTDH3-NtCYP96T6-tTDH1, pTEF1-NtNMT1-tENO2, pCCW12-NtAKR1-tPKG1, pTEF1-OeCPR-tCYC1, pPGK1-CrCYB5-tVPS13. | This study. |
| yMHO57 | yCCD45 | MATa; his3D1; leu2-3_112; ura3-52; trp1-289; pTEF1-SpCas9-tCYC1, pTDH3-NtCYP96T6-tTDH1, pTEF1-NtNMT1-tENO2, pCCW12-NtAKR1-tPKG1, pTEF1-CpCPR-tCYC1, pPGK1-CrCYB5-tVPS13. | This study. |
| yMHO58 | yCCD45 | MATa; his3D1; leu2-3_112; ura3-52; trp1-289; pTEF1-SpCas9-tCYC1, pTDH3-NtCYP96T6-tTDH1, pTEF1-NtNMT1-tENO2, pCCW12-NtAKR1-tPKG1, pTEF1-CrCPR-tCYC1, pPGK1-CrCYB5-tVPS13. | This study. |
| yMHO59 | yCCD45 | MATa; his3D1; leu2-3_112; ura3-52; trp1-289; pTEF1-SpCas9-tCYC1, pTDH3-NtCYP96T6-tTDH1, pTEF1-NtNMT1-tENO2, pCCW12-NtAKR1-tPKG1, pTEF1-AnnCPR-tCYC1, pPGK1-CrCYB5-tVPS13. | This study. |
| yMHO60 | yCCD45 | MATa; his3D1; leu2-3_112; ura3-52; trp1-289; pTEF1-SpCas9-tCYC1, pTDH3-NtCYP96T6-tTDH1, pTEF1-NtNMT1-tENO2, pCCW12-NtAKR1-tPKG1, pTEF1-AtATR1-tCYC1, pPGK1-CrCYB5-tVPS13. | This study. |
| yMHO116 | yCCD45 | MATa; his3D1; leu2-3_112; ura3-52; trp1-289; pTEF1-SpCas9-tCYC1, pTDH3-NtCYP96T6-tTDH1, pTEF1-NtNMT1-tENO2, pCCW12-NtAKR1-tPKG1, pTEF1-AtATR1-tCYC1. | This study. |
| yMHO117 | yCCD45 | MATa; his3D1; leu2-3_112; ura3-52; trp1-289; pTEF1-SpCas9-tCYC1, pTDH3-NtCYP96T6-tTDH1, pTEF1-NtNMT1-tENO2, pCCW12-NtAKR1-tPKG1, pTEF1-AtATR1-tCYC1, pPGK1-NpCYB5-tVPS13. | This study. |
| yMHO118 | yCCD45 | MATa; his3D1; leu2-3_112; ura3-52; trp1-289; pTEF1-SpCas9-tCYC1, pTDH3-NtCYP96T6-tTDH1, pTEF1-NtNMT1-tENO2, | This study. |

| | | | |
|---|---|---|---|
| | | pCCW12-NtAKR1-tPKG1, pTEF1-NpCPR-tCYC1, pPGK1-CrCYB5-tVPS13. | |
| yMHO119 | yCCD45 | MATa; his3D1; leu2-3_112; ura3-52; trp1-289; pTEF1-SpCas9-tCYC1, pTDH3-NtCYP96T6-tTDH1, pTEF1-NtNMT1-tENO2, pCCW12-NtAKR1-tPKG1, pTEF1-NpCPR-tCYC1. | This study. |
| yMHO120 | yCCD45 | MATa; his3D1; leu2-3_112; ura3-52; trp1-289; pTEF1-SpCas9-tCYC1, pTDH3-NtCYP96T6-tTDH1, pTEF1-NtNMT1-tENO2, pCCW12-NtAKR1-tPKG1, pTEF1-NpCPR-tCYC1, pPGK1-NpCYB5-tVPS13. | This study. |
| yMHO121 | yCCD45 | MATa; his3D1; leu2-3_112; ura3-52; trp1-289; pTEF1-SpCas9-tCYC1, pTDH3-NtCYP96T6-tTDH1, pTEF1-NtNMT1-tENO2, pCCW12-NtAKR1-tPKG1, pTEF1-LrCPR2-tCYC1, pPGK1-CrCYB5-tVPS13. | This study. |
| yMHO122 | yCCD45 | MATa; his3D1; leu2-3_112; ura3-52; trp1-289; pTEF1-SpCas9-tCYC1, pTDH3-NtCYP96T6-tTDH1, pTEF1-NtNMT1-tENO2, pCCW12-NtAKR1-tPKG1, pTEF1-LrCPR2-tCYC1. | This study. |
| yMHO123 | yCCD45 | MATa; his3D1; leu2-3_112; ura3-52; trp1-289; pTEF1-SpCas9-tCYC1, pTDH3-NtCYP96T6-tTDH1, pTEF1-NtNMT1-tENO2, pCCW12-NtAKR1-tPKG1, pTEF1-LrCPR2-tCYC1, pPGK1-NpCYB5-tVPS13. | This study. |

#### Supplementary Material References
